# A role for Lin28a in aging-associated decline of adult hippocampal neurogenesis

**DOI:** 10.1101/2022.01.03.474756

**Authors:** Zhechun Hu, Jiao Ma, Huimin Yue, Xiaofang Li, Chao Wang, Liang Wang, Binggui Sun, Zhong Chen, Lang Wang, Yan Gu

## Abstract

Hippocampal neurogenesis declines with aging. Wnt ligands and antagonists within the hippocampal neurogenic niche regulate the proliferation of neural progenitor cells and the development of new neurons, and the changes of their levels in the niche mediate aging-associated decline of neurogenesis. We found that RNA-binding protein Lin28a remained existent in neural progenitor cells and granule neurons in the adult hippocampus, and decreased with aging. Loss of Lin28a inhibited the responsiveness of neural progenitor cells to niche Wnt agonist and reduced neurogenesis, thus impairing pattern separation. Overexpression of Lin28a increased the proliferation of neural progenitor cells, promoted the functional integration of newborn neurons, restored neurogenesis in Wnt-deficient dentate gyrus, and rescued the impaired pattern separation in aging mice. Our data suggest that Lin28a regulates adult hippocampal neurogenesis as an intracellular mechanism by responding to niche Wnt signals, and its decrease is involved in aging-associated decline of hippocampal neurogenesis as well as related cognitive functions.

## Introduction

In the adult mammalian hippocampus, neural stem cells in the subgranular zone (SGZ) of the dentate gyrus (DG) give rise to newborn neurons throughout life (Ming and Song, 2011). Continuously generated newborn neurons integrate into the existing neural circuits, and play essential roles for hippocampal functions, such as hippocampus-dependent mood regulation, memory coding and pattern separation (Anacker and Hen, 2017; Christian et al., 2014; Goncalves et al., 2016).

Accumulating evidence indicates that the proliferative activity of neural progenitor cells (NPCs) and the integration of newborn neurons decrease with aging, resulting in an age-associated decay of neurogenesis (Kuhn et al., 1996; Ngwenya et al., 2015; Spalding et al., 2013), which is believed to be associated with declines in hippocampal-dependent cognitive functions (Seib et al., 2013). Among all the regulating factors, Wnt ligands and antagonists are secreted by local astrocytes and/or neurons in the neurogenic niche, and play essential roles in regulating both the proliferation of NPCs and the development of newborn neurons (Jang et al., 2013; Kuwabara et al., 2009; Lie et al., 2005; Song et al., 2002). Importantly, Wnt signals in the neurogenic niches change dynamically, thus providing a timely regulation of neurogenesis in the adult hippocampus. For instance, Wnt antagonists in the local microenvironment, such as secreted frizzled-related protein 3 (sFRP3) and dickkopf 1 (Dkk1), are important regulators for hippocampal neurogenesis in response to activity and aging (Jang et al., 2013; Seib et al., 2013). However, the intracellular mechanisms underlying Wnt-related regulation of neurogenesis have not been fully revealed.

Lin28 is an RNA-binding protein with a cold-shock domain and a pair of CCHC zinc knuckle RNA-binding domains (Moss et al., 1997). Lin28 has been known to function as a key regulator for self-renewal of embryonic stem cells, and generation of induced pluripotent stem cells (iPSCs) from somatic cells together with Sox2, Oct4 and Nanog (Yu et al., 2007). Recent evidence suggests that Lin28 also plays essential roles in a variety of biological processes, including tissue growth, metabolisms and cancers (Shyh-Chang and Daley, 2013). The two homologs of Lin28 in mammals, Lin28a and Lin28b, are highly expressed during early embryonic development but are thought to be silenced in most adult tissues (Viswanathan and Daley, 2010).

In the developing nervous system, Lin28a has been found to be highly expressed in the NPCs in neural tube during early embryonic stage and at the ventricular/subventricular zone (VZ/SVZ) of the developing cerebral cortex, regulating the generation of neurons and early brain development (Balzer et al., 2010; Yang et al., 2015). Previous studies showed that Lin28a expression dramatically decreases with brain development (Yang et al., 2015). However, recent studies showed that Lin28a remains in NPCs within neurogenic regions in the adult brain and retina (Cimadamore et al., 2013; Yao et al., 2016), raising the possibility that Lin28a may exist and regulate neurogenesis in the adult or even aging brain, which remains elusive.

In this study, we provide the evidence that Lin28a remains in NPCs and neurons in the adult DG and declines with aging. Our results show that Lin28a regulates the proliferation of NPCs and the development of newborn neurons as a downstream intracellular mechanism underlying the regulation of adult neurogenesis by niche Wnt signals. Thus, along with the aging-associated changes of Wnt ligands/inhibitors in the hippocampal neurogenic niche, the decrease of Lin28a in the NPCs serves as one of the intracellular mechanisms mediating the decline of hippocampal neurogenesis. Increasing the expression of Lin28a in hippocampal NPCs in aging mice could rescue their neurogenesis and the ability of pattern separation.

## Results

### Expression of Lin28a in the DG of adult brain decreases with aging

We first set out to examine the presence of Lin28 in the DG, one of the discrete neurogenic regions in the adult mouse brain. We used RNAscope, a high-sensitivity fluorescent in situ hybridization (FISH) method, in brain sections from adult Nestin-GFP mice. Comparing to the signals obtained using negative control (Neg-ctrl) or positive control (Pos-ctrl) probes, Lin28a but not Lin28b mRNA signals exist in the cell body layer of the DG (Figure 1A). Specifically, Lin28a mRNA colocalized with the cell bodies of Nestin^+^ neural stem/progenitor cells, and Prox1^+^ dentate granule cells (DGCs) (Figure 1B).

**Figure 1.**
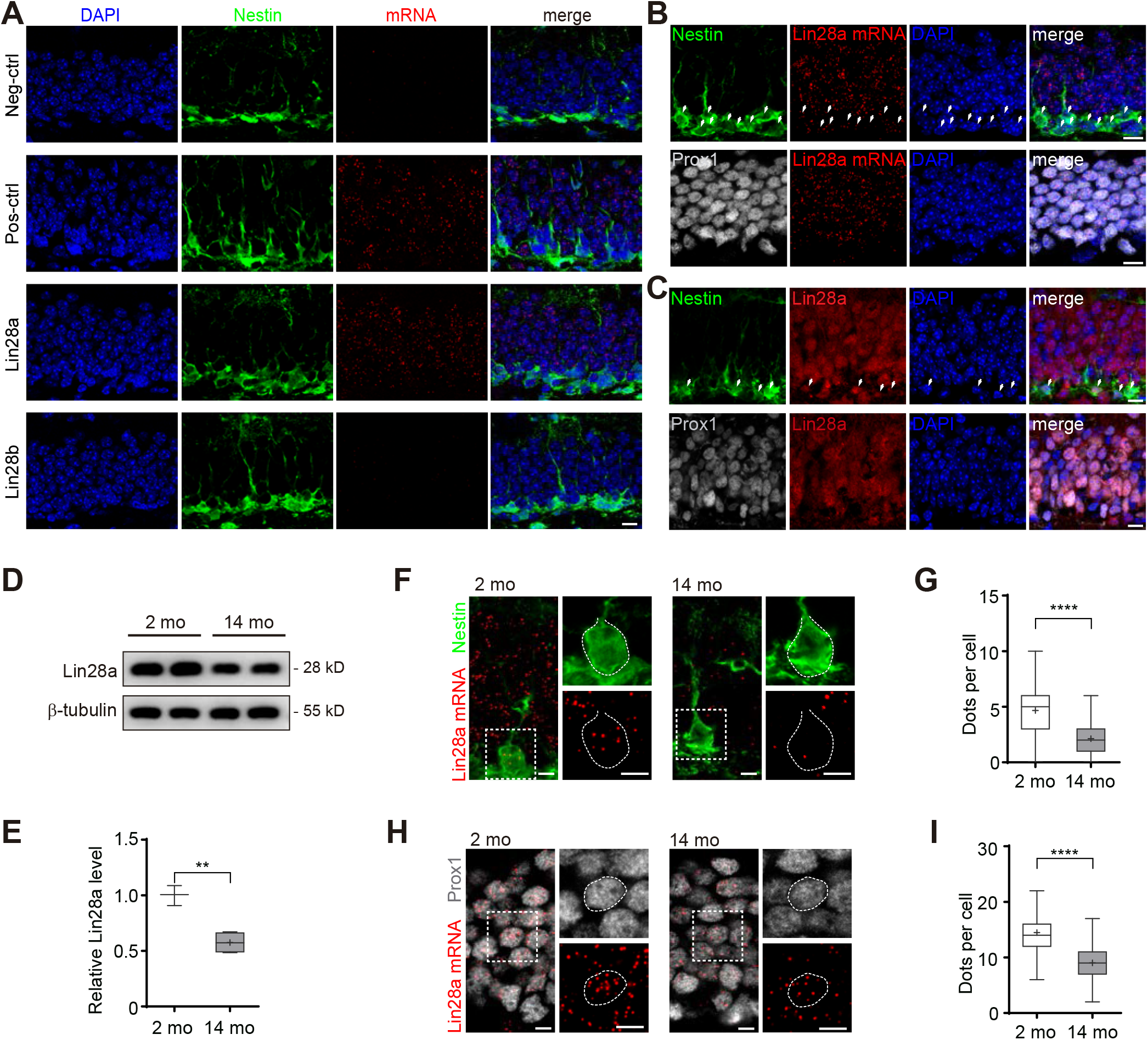
Expression of Lin28a in the DG of adult brain decreases with age. **(A)** Representative RNAscope images obtained using brain sections from Nestin-GFP mice, with negative control (Neg-ctrl), positive control (Pos-ctrl), Lin28a and Lin28b probes, respectively. Scale bar: 20 μm. **(B)** Upper panel: RNAScope images showing Lin28a mRNA in Nestin^+^ NPCs in the DG of adult Nestin-GFP mice. White arrows indicate localization of Lin28a mRNA within cell body of Nestin^+^ NPCs. Scale bar: 10 μm. Lower panel: RNAScope images showing Lin28a mRNA in Prox1^+^ DGCs in adult mice. Scale bar: 10 μm. **(C)** Upper panel: Zoom-in confocal images showing the presence of Lin28a in Nestin^+^ NPCs in the DG of adult Nestin-GFP mice. White arrows indicate colocalization of Lin28a with cell body of Nestin^+^ NPCs. Scale bar: 10 μm. Lower panel: Zoom-in confocal images showing the presence of Lin28a in Prox1^+^ DGCs in adult mice. Scale bar: 10 μm. **(D)** Representative western blotting showing Lin28a expression level in the DG of young adult (2 mo) and aging (14 mo) mice. β-tubulin was used as the internal control. **(E)** Relative expression level of Lin28a in the DG of 2 mo and 14 mo mice. (2 mo: N=3 samples, n=3 mice per sample; 14 mo: N=4 samples, n=3 mice per sample; t_5_=6.162, **P=0.0016). **(F)** RNAScope images showing Lin28a mRNA in Nestin^+^ NPCs in the DG of 2 mo and 14 mo Nestin-GFP mice. Scale bar: 5 μm. **(G)** Number of dots obtained using Lin28a mRNA probes in each Nestin^+^ NPC in the DG of 2 mo and 14 mo mice (2 mo: N=3 mice, n= 82 cells; 14 mo: N=3 mice, n=90 cells; two-tailed unpaired t-test, t_170_=9.427, ****P<0.0001). **(H)** RNAScope images showing Lin28a mRNA in Prox1^+^ DGCs in 2 mo and 14 mo mice. Scale bar: 5 μm. **(I)** Number of dots obtained using Lin28a mRNA probes in each Prox1^+^ DGC in 2 mo and 14 mo mice (2 mo: N=3 mice, n= 99 cells; 14 mo: N=3 mice, n=111 cells; two-tailed unpaired t-test, t_208_=12.21, ****P<0.0001).

To confirm the expression of Lin28a in the adult DG, we next performed immunohistochemistry, and found Lin28a protein exists in the granule cell layer (GCL) of the DG (Figure EV1A). Consistently, Lin28a was observed colocalizing with Nestin^+^ NPCs and Prox1^+^ DGCs (Figure 1C). In contrast, in Nestin-Cre::Lin28a^flox/flox^ mice (Lin28a^f/f^), Lin28a expression was not detected in either the GCL or the SGZ (Figure EV1B).

Interestingly, we found Lin28a protein level in the DG decreased in aging mice (14 months old, 14 mo) compared to young adult mice (2 months old, 2 mo) (Figure 1D and E). Using RNAscope, we were able to analyze Lin28a mRNA level in individual cells, reflected by the number of dots marked by Lin28a mRNA probes in each cell, as previously described (Erben and Buonanno, 2019). We found the number of dots per cell became significantly less in 14 mo animals compared with 2 mo animals, in both Nestin^+^ NPCs (Figure 1F and G) and Prox1^+^ DGCs (Figure 1H and I), indicating an aging-associated decrease of Lin28a expression in these cells.

### Lin28a in hippocampal NPCs are essential for neurogenesis and pattern separation

To find out whether Lin28a decrease in the DG is related to cognitive declines in aging animals, we next trained young adult (2 mo) and aging (10 mo) mice for a series of behavior tests. We found that the 10 mo mice showed decreased novel location recognition (Figure EV2A – C), while the novel object recognition was not affected (Figure EV2D – F). The young and aging animals didn’t show significant difference in single-trial contextual fear memory test (Figure EV2G and H). However, when we tested the animals with a pattern separation behavior paradigm (Figure 2A), as previously reported (Sahay et al., 2011a), we found that the 10 mo mice discriminated the two similar contexts A and B in the 6th test session, much later than the 2 mo mice, which showed contextual discrimination in the 4th session (Figure 2B). This result suggested that aging is associated with a decreased ability of pattern separation, consistent with previous studies on aging human and non-human primate subjects (Dillon et al., 2017; Ngwenya et al., 2015).

**Figure 2.**
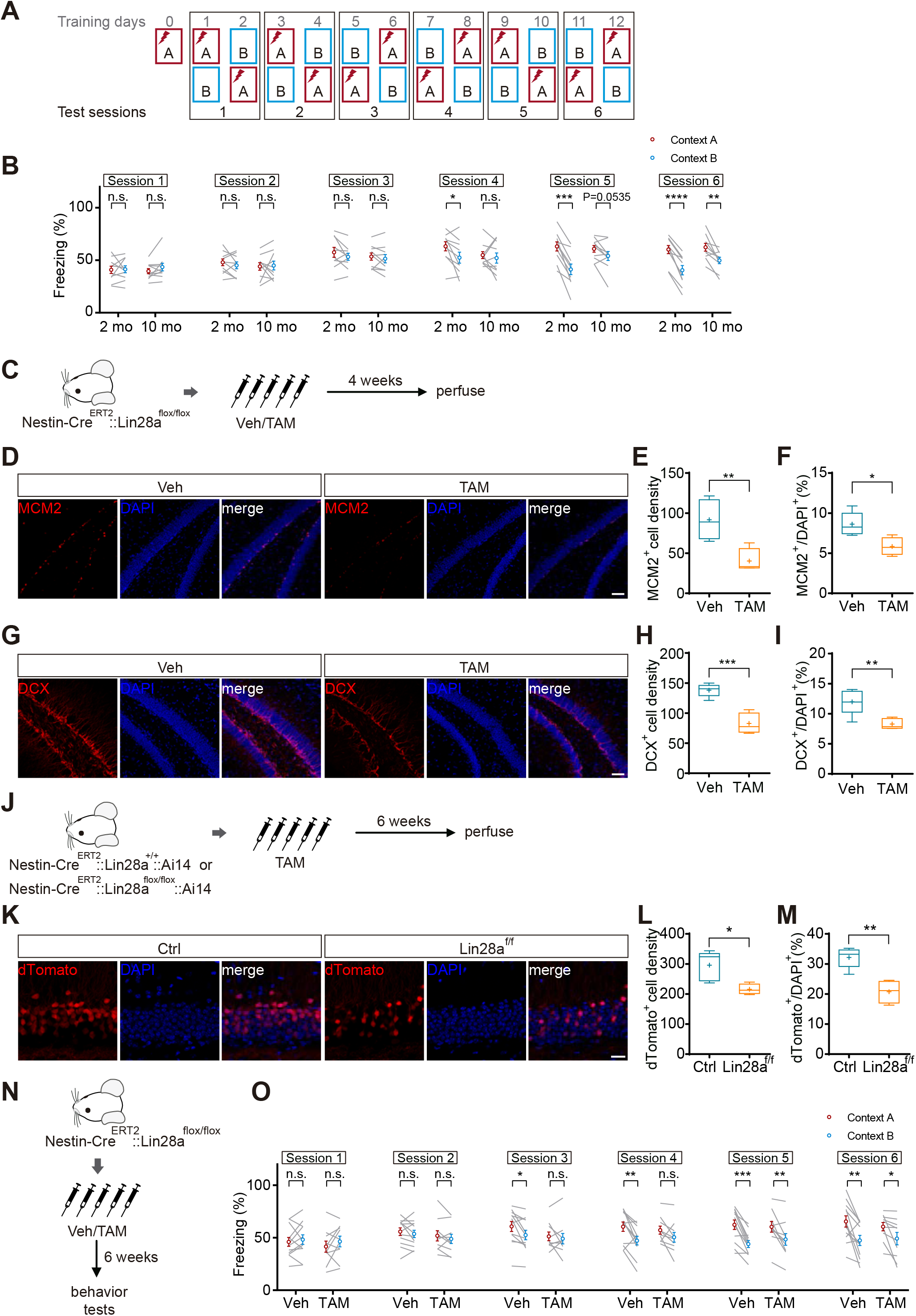
Lin28a is necessary for adult hippocampal neurogenesis and pattern separation. **(A)** Experimental paradigm for testing pattern separation in mice. **(B)** The freezing of 2 mo and 10 mo mice in contexts A and B in test sessions 1 through 6. Young mice (2 mo) discriminated the two contexts in session 4, while aging mice (10 mo) showed discrimination in session 6. (2 mo mice n=10, 10 mo mice n=10; two-tailed paired t-test; session 1: 2 mo t_9_=0.1441, P=0.8886; 10 mo t_9_=1.609, P=0.1421; session 2: 2 mo t_9_=1.036, P=0.3272; 10 mo t_9_=0.2128, P=0.8362; session 3: 2 mo t_9_=1.391, P=0.1976; 10 mo t_9_=0.8787, P=0.4024; session 4: 2 mo t_9_=2.672, *P=0.0255; 10 mo t_9_=0.6441, P= 0.5356; session 5: 2 mo t_9_=6.572, ***P= 0.0001; 10 mo t_9_=2.220, P=0.0535; session 6: 2 mo t_9_=7.818, ****P<0.0001; 10 mo t_9_=3.897, **P=0.0036). **(C)** Diagram showing TAM or Veh was administered to Nestin-Cre^ERT2^::Lin28a^flox/flox^ mice to knockout Lin28a from Nestin^+^ NPCs. Mice were perfused four weeks later for examination. **(D)** Confocal images showing MCM2^+^ proliferating cells in the SGZ of Nestin-Cre^ERT2^::Lin28a^flox/flox^ mice treated with Veh or TAM. Scale bar: 50 μm. **(E)** Knocking out Lin28a from Nestin^+^ NPCs decreased the density of MCM2^+^ cells in the SGZ (Veh n=5 mice, TAM n=4 mice; two-tailed unpaired t-test, t_7_=3.638, **P=0.0083). **(F)** Knocking out Lin28a from Nestin^+^ NPCs decreased the percentage of MCM2^+^ cells in total cells (DAPI^+^) in the SGZ and GCL (Veh n=5 mice, TAM n=4 mice; two-tailed unpaired t-test, t_7_=3.116, *P=0.0158). **(G)** Confocal images showing DCX^+^ cells in the DG of Nestin-Cre^ERT2^::Lin28a^flox/flox^ mice treated with Veh or TAM. Scale bar: 50 μm. **(H)** Loss of Lin28a from Nestin^+^ NPCs decreased the density of DCX^+^ cells in the GCL (Veh n=5 mice, TAM n=5 mice; two-tailed unpaired t-test, t_8_=6.252, ***P=0.00023). **(I)** Loss of Lin28a from Nestin^+^ NPCs decreased the percentage of DCX^+^ cells in total cells (DAPI^+^) in the SGZ and GCL (Veh n=5 mice, TAM n=5 mice; two-tailed unpaired t-test, t_8_=3.656, **P=0.0064). **(J)** Diagram showing TAM was administered to Nestin-Cre^ERT2^::Lin28a^+/+^::Ai14 or Nestin-Cre^ERT2^::Lin28a^flox/flox^::Ai14 mice. Mice were perfused six weeks later for examination. **(K)** Confocal images showing dTomato^+^ cells in the GCL of Nestin-Cre^ERT2^:: Lin28a^+/+^::Ai14 (Ctrl) or Nestin-Cre^ERT2^::Lin28a^flox/flox^::Ai14 (Lin28a^f/f^) mice. Scale bar: 20 μm. **(L)** Loss of Lin28a from Nestin^+^ NPCs decreased the density of dTomato^+^ cells in the GCL (Ctrl n=5 mice, Lin28a^f/f^ n=4 mice; two-tailed unpaired t-test, t_7_=3.113, *P=0.0170). **(M)** Loss of Lin28a from Nestin^+^ NPCs decreased the percentage of dTomato^+^ cells in total cells (DAPI^+^) in the GCL (Ctrl n=5 mice, Lin28a^f/f^ n=4 mice; two-tailed unpaired t-test, t_7_=4.780, **P=0.0020). **(N)** Schematics showing that Nestin-Cre^ERT2^::Lin28a^flox/flox^ mice were treated with Veh (Ctrl) or TAM (Lin28a^f/f^), followed by behavioral tests six weeks later. **(O)** The freezing of Veh- and TAM-treated mice in contexts A and B in test sessions 1 through 6. (Veh mice n=12, TAM mice n=10; two-tailed paired t-test; session 1: Veh t_11_=0.6717, P=0.5156; TAM t_9_=1.151, P=0.2795; session 2: Veh t_11_=0.8907, P=0.3921; TAM t_9_=1.177, P=0.2694; session 3: Veh t_11_=2.867, *P=0.0153; TAM t_9_=0.6078, P=0.5583; session 4: Veh t_11_=3.427, **P= 0.0057; TAM t_9_=1.925, P=0.0863; session 5: Veh t_11_=5.504, ***P= 0.0002; TAM t_9_=3.434, **P=0.0075; session 6: Veh t_11_=3.530, **P=0.0047; TAM t_9_=2.614, *P=0.0281).

Because pattern separation is believed to be one of the hippocampal functions most related to neurogenesis (Sahay et al., 2011b), we checked the level of neurogenesis in the DG of the 2 mo and 10 mo mice. Indeed, 10 mo mice showed significantly less doublecortin (DCX)-positive cells in the DG (Figure EV2I and J), indicating an aging-associated decline of hippocampal neurogenesis, consistent with previous reports (Seib et al., 2013).

To test whether Lin28a in the hippocampal NPCs is related to adult neurogenesis, we then knocked out Lin28a from Nestin^+^ NPCs in Nestin-Cre^ERT2^::Lin28a^flox/flox^ mice by tamoxifen (TAM) treatment, and examined the neurogenesis in the DG of these mice four weeks later (Figure 2C). We found loss of Lin28a from Nestin^+^ NPCs significantly decreased the number of cells expressing proliferation marker minichromosome maintenance complex component 2 (MCM2) (Figure 2D), as indicated by the density of MCM2^+^ cells in the SGZ and the percentage of MCM2^+^ cells in total DAPI^+^ cells in the GCL (Figure 2E and F). Similarly, DCX-positive cells significantly decreased in the DG of the TAM-treated animals, compared to the mice treated with vehicle (Veh) (Figure 2G – I). To further verify whether loss of Lin28a in NPCs indeed results in fewer adult-born neurons, we treated and Nestin-Cre^ERT2^::Lin28a^flox/flox^::Ai14 (Lin28a^f/f^) mice and Nestin-Cre^ERT2^:: Lin28a^+/+^::Ai14 (Ctrl) mice of the same age and genetic background with tamoxifen, and examined the number of dTomato-labeled cells in the DG six weeks later (Figure 2J). We found that lack of Lin28a indeed decreased the density and the proportion of dTomato^+^ cells in the DG of Lin28a^f/f^ mice, compared with Ctrl mice (Figure 2K – M). These data suggest that Lin28a is necessary for the generation of new neurons from adult hippocampal NPCs, whereas loss of Lin28a leads to decreased neurogenesis in the adult DG.

We then tested Nestin-Cre^ERT2^::Lin28a^flox/flox^ mice for pattern separation, six weeks after Veh or TAM treatment (Figure 2N). In consistence with the decreased hippocampal neurogenesis, TAM-treated mice discriminated the two similar contexts A and B later than Veh-treated mice did (Figure 2O), mimicking the decreased ability of pattern separation in aging animals.

In addition, similar to the behavioral phenotypes seen in the aging animals, the TAM-treated Nestin-Cre^ERT2^::Lin28a^flox/flox^ mice didn’t show significant difference in single-trial contextual fear memory test (Figure EV3A and B), or novel object recognition (Figure EV3C – E). However, TAM-treated mice showed impaired novel location recognition (Figure EV3F – H).

These results suggest that the decreased Lin28a level in hippocampal NPCs may contribute to the decline of neurogenesis and the ability of pattern separation in the aging animals.

### Overexpression of Lin28a increases the progenies of NPCs in the adult DG

Next, to investigate how Lin28a regulates neural progenitors in the adult brain, we generated FLEX retroviral vectors that express GFP (Ctrl) or GFP-p2A-Lin28a (Lin28a-OE) (Figure 3A). We injected diluted virus into the DG of adult Nestin-Cre mice, so that GFP or Lin28a could be expressed upon the presence of Cre recombinase in individual dividing Nestin^+^ NPCs and their progenies in a sparse distribution (Figure 3B). With this method, we were able to trace the progenies from individual active neural stem cells, as previously described (Kirschen et al., 2017). We examined GFP labeled cell clusters 7 days after viral injection, and found that the overexpression of Lin28a significantly increased the size of cell clusters generated from individual NPCs, from an average of 1.77±0.11 to 3.23±0.26 cells per cluster (Figure 3C – E).

**Figure 3.**
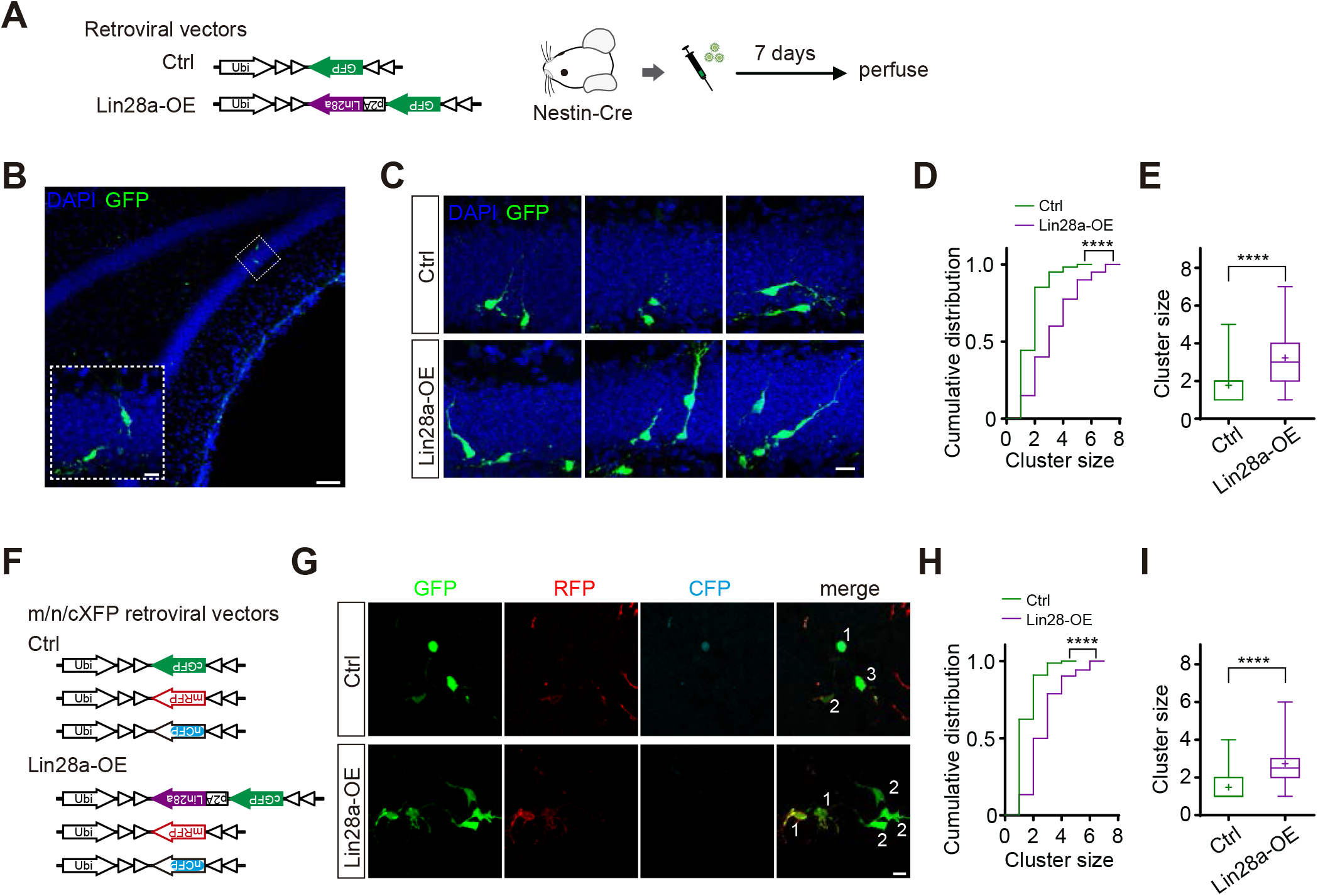
Overexpression of Lin28a increases the progenies of individual NPCs. **(A)** Schematics showing that FLEX retroviral vectors expressing GFP (Ctrl) or GFP-p2A-Lin28a (Lin28a-OE) were injected into the DG of Nestin-Cre mice. **(B)** A confocal image showing the sparse labeling of an individual cluster of newborn cells in the DG. Scale bar: 50 μm. Inset shows an enlarged image of the 2-cell cluster in the square area marked by dotted line. Scale bar: 10 μm. **(C)** Confocal images showing examples of cell clusters labeled by Ctrl or Lin28a-OE retroviruses in the DG of Nestin-Cre mice. Scale bar: 10 μm. **(D)** Cumulative distribution of the size of cell clusters labeled by Ctrl or Lin28a-OE retroviruses. (Ctrl N=11 mice, n=61 cell clusters; Lin28a-OE N=8 mice, n=40 cell clusters; Kolmogorov-Smirnov test, ****P<0.0001). **(E)** The average size of cell clusters labeled by Ctrl or Lin28a-OE retroviruses. (Ctrl N=11 mice, n=61 cell clusters; Lin28a-OE N=8 mice, n=40 cell clusters; two-tailed unpaired t-test, t_99_=5.700, ****P<0.0001). **(F)** Schematics showing combination of Cre-dependent m/n/cXFP retroviral vectors used for clonal labeling in Nestin-Cre mice in Ctrl or Lin28a-OE groups. **(G)** Confocal images showing labeling of cell clusters in the DG of Nestin-Cre mice by Ctrl or Lin28a-OE combination of m/n/cXFP retroviral vectors. Different numbers indicate different cell clusters. In Lin28a-OE group, only GFP^+^ cell clusters were selected for analysis. Scale bar: 10 μm. **(H)** Cumulative distribution of the size of cell clusters labeled in Ctrl or Lin28a-OE groups. (Ctrl N=8 mice, n=77 cell clusters; Lin28a-OE N=7 mice, n=52 cell clusters; Kolmogorov-Smirnov test, ****P<0.0001). **(I)** The average size of cell clusters labeled by Ctrl or Lin28a-OE combination of m/n/cXFP retroviral vectors. (Ctrl N=8 mice, n=77 cell clusters; Lin28a-OE N=7 mice, n=52 cell clusters; two-tailed unpaired t-test, t_127_=7.123, ****P<0.0001).

To confirm this result and avoid possible mixture of adjacently-labeled cell clusters, we designed a FLEX m/n/cXFP retroviral reporter system that expresses membranous RFP (mRFP), nuclear CFP (nCFP) or cytosolic GFP (cGFP) in random combinations depending on Cre recombinase expression (Figure 3F). Seven days post viral injection, we analyzed labeled cell clusters in both Ctrl and Lin28a-OE groups (Figure 3G). Consistent with the data shown in Figure 3D and E, cells overexpressing Lin28a (containing cGFP) exhibited increased cluster size of 2.73±0.18 cells, compared to 1.48±0.08 cells in Ctrl clusters (Figure 3H and I), confirming that the overexpression of Lin28a indeed increased the number of cells generated from each individual NPC.

### The effect of Lin28a overexpression on cell cycle, survival and differentiation

To investigate how Lin28a regulate the number of progenies of NPCs, we next tested whether Lin28a could regulate the cell cycle of hippocampal NPCs. Two days after the injection of FLEX retroviruses (Figure 3A) into the DG of Nestin-Cre mice, we started treating the animals with BrdU every 6 hours for 5 times. Seven days after viral injection, we perfused the animals and stained the brain sections for BrdU and MCM2 (Figure 4A). By this method, we could label dividing Nestin^+^ NPCs with GFP at their first division, and then mark their second division with BrdU if the mitosis happened during the window of BrdU administration, while a third division could be detected by MCM2 staining (Figure 4B). We found overexpressing Lin28a in NPCs significantly increased the percentage of GFP^+^BrdU^+^ and GFP^+^BrdU^+^MCM2^+^ cells in GFP-labeled cells (Figure 4C), indicating that the overexpression of Lin28a enlarges the size of cell clusters via increasing the number of cell cycles of NPCs.

**Figure 4.**
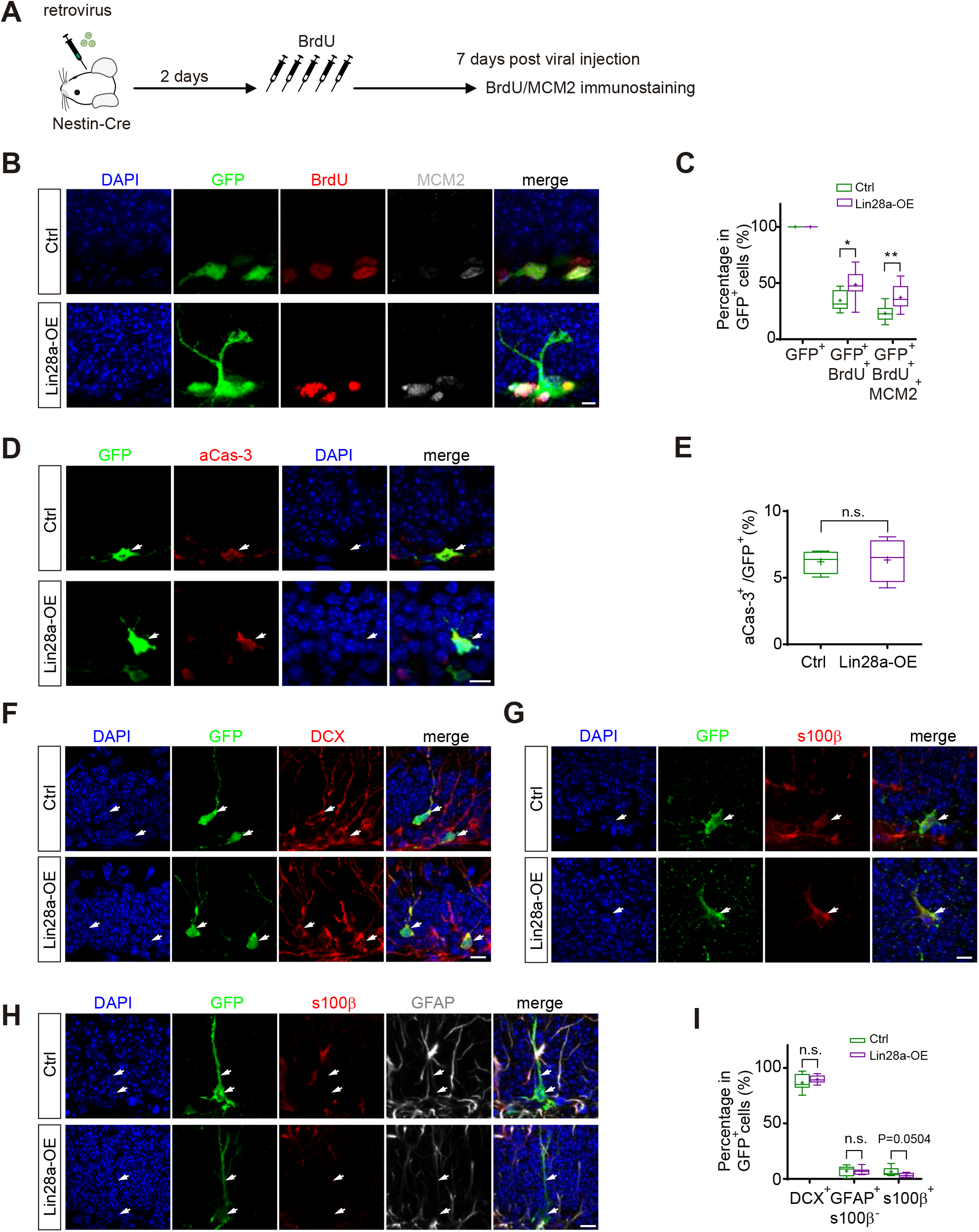
The effect of Lin28a overexpression on cell cycle, survival and differentiation. **(A)** Experimental scheme showing the analysis of cell cycles of hippocampal NPCs *in vivo*. **(B)** Confocal images showing GFP-labeled newborn cells in the DG using Ctrl or Lin28a-OE retroviruses, co-labeled with BrdU and/or MCM2. Scale bar: 5 μm. **(C)** Percentage of GFP^+^BrdU^+^ cells (two-tailed unpaired t-test, t_16_=2.7507, *P=0.0142) and GFP^+^BrdU^+^MCM2^+^ cells (two-tailed unpaired t-test, t_16_=3.5555, **P=0.0040) in GFP-labeled cells (Ctrl N=10 mice, n=290 cells; Lin28a-OE N=8 mice, n=253 cells). **(D)** Images showing aCas-3^+^ GFP-expressing newborn cells labeled by Ctrl and Lin28a-OE retroviruses in the DG. White arrows indicate an aCas-3^+^ GFP-labeled cell. Scale bar: 10 μm. **(E)** Percentage of aCas-3^+^ cells in GFP-labeled newborn cells in the DG. (Ctrl N=4 mice, n=794 cells; Lin28a-OE N=4 mice, n=339 cells; two-tailed unpaired t-test, t_6_=0.1449, P=0.8896). **(F)** Images showing DCX^+^ GFP-expressing newborn cells labeled by Ctrl or Lin28a-OE retroviruses in the DG. White arrows indicate GFP^+^DCX^+^ cells. Scale bar: 10 μm. **(G)** Images showing a GFP^+^s100β^+^ astrocytes labeled by Ctrl or Lin28a-OE retroviruses in the DG, indicated by white arrows. Scale bar: 10 μm. **(H)** Images showing GFP^+^GFAP^+^s100β^-^ radial glia-like NPCs labeled by Ctrl or Lin28a-OE retroviruses in the DG, indicated by white arrows. Scale bar: 10 μm. **(I)** Percentage of DCX^+^ (two-tailed unpaired t-test, t_16_=1.2006, P=0.2474), GFAP^+^s100β^-^ (two-tailed unpaired t-test, t_16_=0.1215, P=0.9048) and s100β^+^ (two-tailed unpaired t-test, t_16_=2.1156, P=0.0504) cells in GFP-labeled newborn cells (Ctrl N=10 mice; Lin28a-OE N=8 mice), seven days after viral injection.

Next, to test whether the change in the survival of newborn cells could also count for the increase of cluster size, we stained for activated caspase-3 (aCas-3) four days after retroviral injection (Figure 4D). We found colocalization of aCas-3 with GFP-labeled newborn cells did not show significant difference between cells overexpressing Lin28a and Ctrl cells expressing GFP only (Figure 4E).

To further find out whether Lin28a controls the differentiation of newborn cells, we overexpressed Lin28a in Nestin^+^ NPCs and quantified proportion of DCX^+^ newborn neurons (Figure 4F), s100β^+^ astrocytes (Figure 4G) and GFAP^+^s100β^-^ neural stem/progenitor cells (Figure 4H) in GFP-labeled newborn cells, seven days after viral injection. We found the overexpression of Lin28a did not significantly change the proportion of GFAP^+^s100β^-^ cells, but slightly increased the proportion of DCX^+^ cells and decreased the percentage of s100β^+^ cells (Figure 4I). This result suggests that Lin28a may potentially support the newborn cells to differentiate towards neuronal lineage, and inhibit the differentiation of newborn cells towards astrocytes.

### Lin28a regulates the development and functional integration of new neurons in the adult DG

Since Lin28a is also detected in DGCs, we wondered whether Lin28a regulates the development and functional integration of the newborn neurons. We then labeled the newborn neurons in the DG of adult C57BL/6 mice using retroviral vectors expressing GFP (Ctrl) or GFP-p2A-Lin28a (Lin28a-OE), and examined the morphological development of labeled newborn neurons at 1, 2, 3 and 4 weeks post retroviral injection (wpi) (Figure 5A). Our data showed that neurons overexpressing Lin28a exhibited significantly more complex dendrites (Figure 5B), longer total dendritic length (Figure 5C) and more dendritic branches at all developmental stages (Figure 5D), suggesting that the overexpression of Lin28a facilitates the development of newborn neurons.

**Figure 5.**
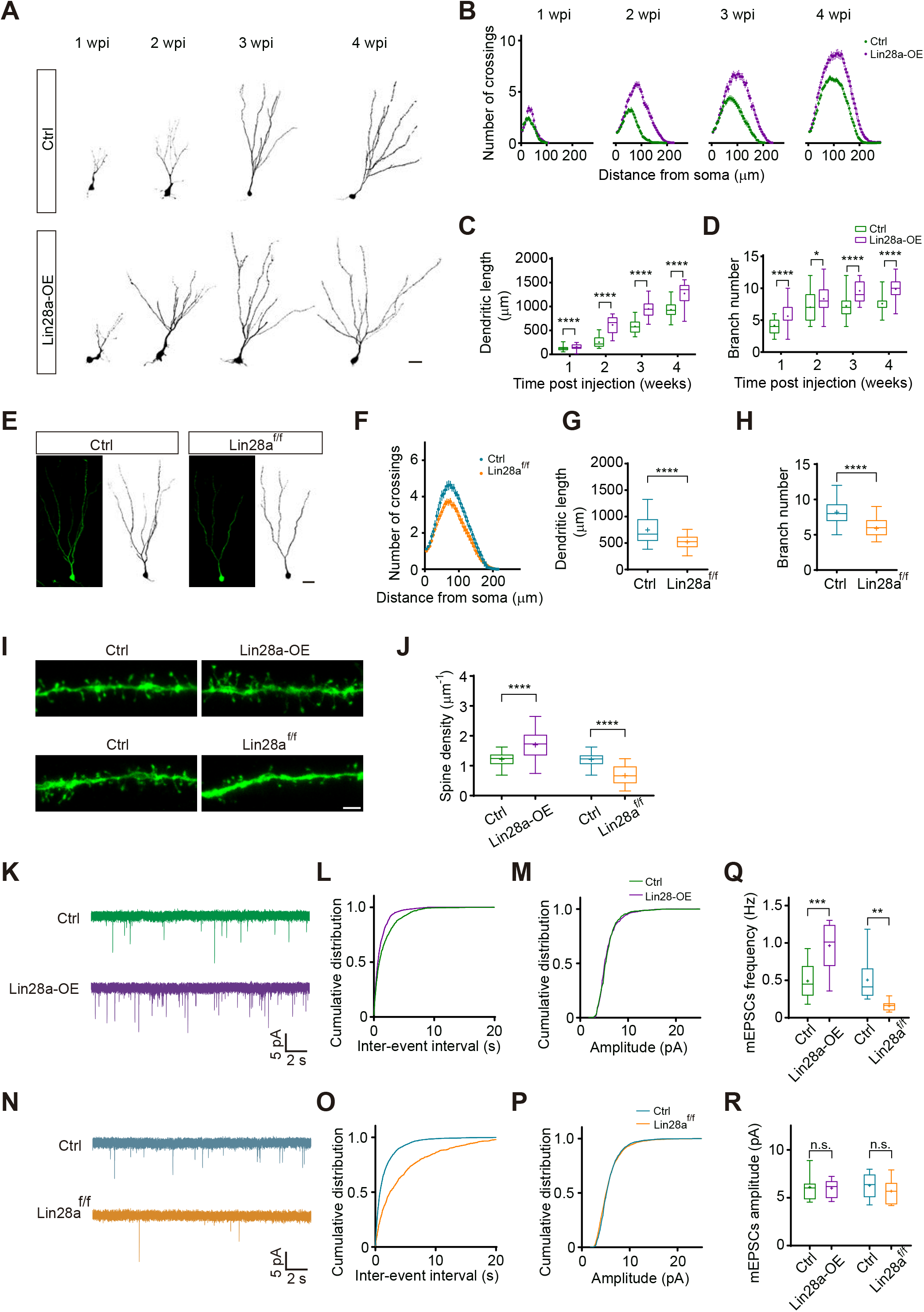
Lin28a regulates the development and functional integration of newborn neurons in the adult DG. **(A)** Representative images of newborn neurons in the DG of adult mice labeled by retrovirus expressing GFP (Ctrl) or GFP-p2A-Lin28a (Lin28a-OE) at 1, 2, 3, and 4 wpi. Images were converted into black and white. Scale bar: 20 μm. **(B)** Sholl analysis of dendrites of Ctrl or Lin28a-OE newborn neurons at 1, 2, 3, and 4 wpi. **(C)** Lin28a-OE neurons showed longer total dendritic length than Ctrl neurons at 1, 2, 3, and 4 wpi. (1 wpi: Ctrl N=4 mice, n=60 cells; Lin28a-OE N=4 mice, n=52 cells; two-tailed unpaired t-test, t_110_=4.742, ****P<0.0001; 2 wpi: Ctrl N=3 mice, n=27 cells; Lin28a-OE N=3 mice, n=44 cells; two-tailed unpaired t-test, t_69_=9.921, ****P<0.0001; 3 wpi: Ctrl N=3 mice, n=34 cells; Lin28a-OE N=3 mice, n=23 cells; two-tailed unpaired t-test, t_55_=8.371, ****P<0.0001; 4 wpi: Ctrl N=3 mice, n=47 cells; Lin28a-OE N=3 mice, n=23 cells; two-tailed unpaired t-test, t_68_=7.6342, ****P<0.0001). **(D)** Lin28a-OE neurons showed more dendritic branches than Ctrl neurons at 1, 2, 3, and 4 wpi. (1 wpi: Ctrl N=4 mice, n=60 cells; Lin28a-OE N=4 mice, n=52 cells; two-tailed unpaired t-test, t_110_=5.263, ****P<0.0001; 2 wpi: Ctrl N=3 mice, n=27 cells; Lin28a-OE N=3 mice, n=44 cells; two-tailed unpaired t-test, t_69_=2.319, *P=0.0233; 3 wpi: Ctrl N=3 mice, n=34 cells; Lin28a-OE N=3 mice, n=23 cells; two-tailed unpaired t-test, t_55_=4.977, ****P<0.0001; 4 wpi: Ctrl N=3 mice, n=47 cells; Lin28a-OE N=3 mice, n=23 cells; two-tailed unpaired t-test, t_68_=6.619, ****P<0.0001). **(E)** Images of newborn neurons in Lin28a^flox/flox^ mice labeled by retrovirus expressing GFP (Ctrl) or GFP-p2A-Cre (Lin28a^f/f^) at 3 wpi. Scale bar: 20 μm. **(F)** Sholl analysis of dendrites of Ctrl or Lin28a^f/f^ newborn neurons at 3 wpi. **(G)** Lin28a^f/f^ neurons showed shorter total dendritic length than Ctrl neurons at 3 wpi. (Ctrl N=3 mice, n=26 cells; Lin28a^f/f^ N=4 mice, n=53 cells; two-tailed unpaired t-test, t_77_=5.179, ****P<0.0001). **(H)** Lin28a^f/f^ neurons showed fewer dendritic branches than Ctrl neurons at 3 wpi. (Ctrl N=3 mice, n=26 cells; Lin28a^f/f^ N=4 mice, n=53 cells; two-tailed unpaired t-test, t_77_=6.830, ****P<0.0001). **(I)** Confocal images showing dendritic spines of Ctrl or Lin28a-OE newborn neurons in C57 mice (upper panel), and Ctrl or Lin28a^f/f^ newborn neurons in Lin28a^flox/flox^ mice (lower panel) at 3 wpi. Scale bar: 2 μm. **(J)** Spine density in newborn neurons at 3 wpi. (C57 mice: Ctrl N=3 mice, n=30 dendritic segments; Lin28-OE N=3 mice, n=30 dendritic segments; two-tailed unpaired t-test, t_58_=5.429, P<0.0001; Lin28a^flox/flox^ mice: Ctrl N=3 mice, n=22 dendritic segments; Lin28^f/f^ N=3 mice, n=18 dendritic segments; two-tailed unpaired t-test, t_38_=6.248, ****P<0.0001). **(K)** Representative traces of mEPSCs recorded from Ctrl and Lin28a-OE newborn neurons at 3 wpi. **(L)** Cumulative distribution of inter-event intervals of mEPSCs in Ctrl and Lin28a-OE neurons. **(M)** Cumulative distribution of amplitude of mEPSCs in Ctrl and Lin28a-OE neurons. **(N)** Representative traces of mEPSCs recorded from Ctrl and Lin28a^f/f^ neurons at 3 wpi. **(O)** Cumulative distribution of inter-event intervals of mEPSCs in Ctrl and Lin28a^f/f^ neurons. **(P)** Cumulative distribution of amplitude of mEPSCs in Ctrl and Lin28a^f/f^ neurons. **(Q)** mEPSCs frequency increased in Lin28a-OE neurons but decreased in Lin28a^f/f^ neurons. (C57 mice: Ctrl N=3 mice, n=12 neurons; Lin28-OE N=3 mice, n=12 neurons; two-tailed unpaired t-test, t_22_=4.301, ***P=0.0003; Lin28a^flox/flox^ mice: Ctrl N=3 mice, n=9 neurons; Lin28a^f/f^ N=3 mice, n=10 neurons; two-tailed unpaired t-test, t_17_=3.509, **P=0.0027). **(R)** mEPSCs amplitude did not significantly change in Lin28a-OE or Lin28a^f/f^ neurons. (C57 mice: Ctrl N=3 mice, n=12 neurons; Lin28-OE N=3 mice, n=12 neurons; two-tailed unpaired t-test, t_22_=0.2229, P=0.8257; Lin28a^flox/flox^ mice: Ctrl N=3 mice, n=9 neurons; Lin28^f/f^ N=3 mice, n=10 neurons; two-tailed unpaired t-test, t_17_=1.042, P=0.3119).

To test whether Lin28a is necessary for the development of newborn neurons, we injected retrovirus expressing GFP-p2A-Cre into the DG of Lin28a^flox/flox^ mice to knockout Lin28a from labeled newborn neurons (Lin28a^f/f^), while Ctrl cells were labeled by retrovirus expressing GFP only (Figure 5E). At 3 wpi, we found Lin28a^f/f^ newborn neurons showed less complex dendrites (Figure 5F), shorter dendritic length (Figure 5G) and less dendritic arborizations when compared with Ctrl cells (Figure 5H).

Since the third week is the critical time period during which newborn DGCs form synaptic connections with afferent glutamatergic projections (Kumamoto et al., 2012), we further analyzed the dendritic spines of the newborn neurons at 3 wpi. We found that Lin28a-OE newborn neurons exhibited greatly increased spine density, while Lin28a^f/f^ newborn neurons showed significantly less dendritic spines than Ctrl cells expressing GFP only (Figure 5I and J). To verify whether Lin28a regulates the functional synaptic integration of newborn neurons, we then did whole-cell patch-clamp recordings on labeled newborn neurons and recorded the miniature excitatory postsynaptic currents (mEPSCs) at 3 wpi. We found that the overexpression of Lin28a in newborn neurons significantly increased mEPSCs frequency without affecting mEPSCs amplitude (Figure 5K – M), whereas knocking out Lin28a from newborn neurons significantly reduced mEPSCs frequency but not amplitude (Figure 5N – R). Thus, these data suggest that Lin28a facilitated, whereas loss of Lin28a hindered, the development and functional synaptic integration of newborn neurons in the adult brain.

### Lin28a is involved in Wnt-mediated regulation of neurogenesis

Given Wnt signaling regulates age-dependent decline of neurogenesis in the adult hippocampus (Seib et al., 2013), and β-catenin promotes the expression of Lin28a in a variety of cell types (Cai et al., 2013; Yao et al., 2016), we wondered whether Lin28a is the intracellular mediator through which Wnt signaling regulates adult hippocampal neurogenesis. We injected lentivirus expressing Wnt3a, a Wnt agonist expressed in hippocampal neurogenic niche (Kuwabara et al., 2009; Lie et al., 2005), into the DG of adult mice (Figure 6A – C). We found an increased level of β-catenin in the DG (Figure 6D and E), suggesting an enhanced canonical Wnt signaling. Using RNAscope, we found the amount of Lin28a mRNA in Sox2-expressing NPCs significantly increased in animals injected with Wnt3a-expressing virus, compared to those injected with control virus expressing GFP only (Figure 6F and G). These data suggest that the expression of Lin28a in NPCs was increased by enhanced Wnt signaling.

**Figure 6.**
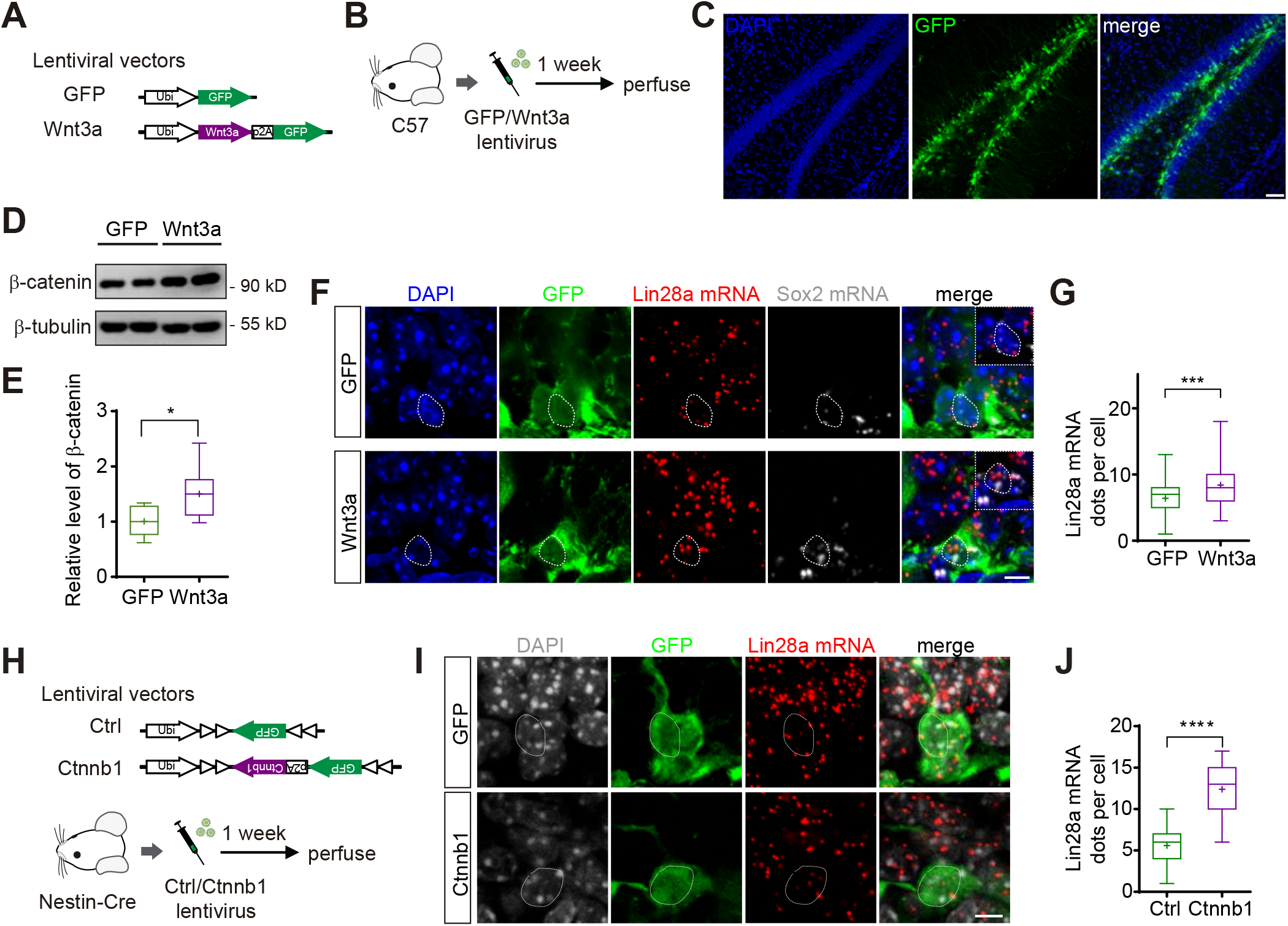
Lin28a expression in NPCs is regulated by Wnt-β-catenin signaling. **(A)** Schematics showing lentiviral vectors expressing GFP or Wnt3a. **(B)** Schematics showing that lentiviral vectors were injected into the DG of adult C57 mice. Mice were perfused for examination one week later. **(C)** Confocal image showing infection of cells in the DG by lentivirus expressing Wnt3a-p2A-GFP. Scale bar: 50 μm. **(D)** Representative western blotting showing β-catenin expression level in the DG of animals injected with lentiviruses expressing GFP or Wnt3a. β-tubulin was used as internal control. **(E)** Relative expression level of β-catenin in the DG of animals injected with GFP or Wnt3a lentiviruses (GFP: n=8 mice per sample; Wnt3a: n=9 mice per sample; t_15_=2.7375, *P=0.0153). **(F)** RNAScope images showing colocalization of Lin28a mRNA with cell bodies of Sox2-expressing NPCs (indicated by Sox2 mRNA) in animals injected with GFP or Wnt3a lentiviruses. Dotted lines outline the cell bodies of interested cells. Insets in the merged images show the localization of Lin28a and Sox2 mRNA around the nucleus (DAPI) within the cell bodies. Scale bar: 5 μm. **(G)** Wnt3a increased the mRNA expression level of Lin28a in Sox2-expressing NPCs. (GFP: N=3 mice, n= 56 cells; Wnt3a: N=3 mice, n=62 cells; two-tailed unpaired t-test, t_116_=3.9875, ***P=0.0001). **(H)** Upper panel: Schematics showing FLEX lentiviral vectors expressing GFP (Ctrl) or GFP-p2A-Ctnnb1 (Ctnnb1). Lower panel: Schematics showing that FLEX lentiviral vectors expressing GFP or GFP-p2A-Ctnnb1 were injected into the DG of adult Nestin-Cre mice. Mice were perfused for examination one week later. **(I)** RNAScope images showing colocalization of Lin28a mRNA with cell bodies of GFP-labeled NPCs in the DG of Nestin-Cre mice injected with Cre-dependent lentiviral vectors expressing GFP or Ctnnb1. Scale bar: 5 μm. **(J)** Upregulation of Ctnnb1 increased the mRNA expression level of Lin28a in NPCs. (GFP: N=3 mice, n= 63 cells; Ctnnb1: N=3 mice, n=37 cells; two-tailed unpaired t-test, t_98_=13.57, ****P<0.0001).

To further confirm whether upregulation of β-catenin could directly enhance the expression of Lin28a in NPCs, we constructed a FLEX lentiviral vector expressing mouse Ctnnb1, which codes the protein catenin beta 1 (Figure 6H). We injected the virus into the DG of Nestin-Cre mice, and performed RNAscope a week later. We found overexpressing Ctnnb1 in Nestin^+^ NPCs indeed increased the mRNA level of Lin28a in these labeled cells (Figure 6I and J).

Next, we injected lentivirus expressing Wnt3a into the DG of Nestin-Cre^ERT2^::Lin28a^flox/flox^ mice, one week after TAM administration (Figure 7A), by which Lin28a was knocked out from Nestin^+^ NPCs (Lin28a^f/f^). Four more weeks later, we found Wnt3a lentivirus significantly increased DCX^+^ cells in the DG of Cre^−/−^ littermates (Ctrl). In contrast, upregulation of Wnt3a was not able to increase the number of DCX^+^ cells in the DG of Lin28a^f/f^ animals (Figure 7B – D). To confirm this finding, we next used Nestin-Cre^ERT2^::Lin28a^flox/flox^::Ai14 mice to label the progenies of the NPCs, from which Lin28a was knocked out (Lin28a^f/f^); Nestin-Cre^ERT2^::Lin28a^+/+^::Ai14 mice of the same age were used as control (Ctrl) (Figure 7E). Consistently, we found Wnt3a lentivirus significantly increased dTomato^+^ cells in the DG of control mice. Whereas, upregulation of Wnt3a could not increase the number of dTomato^+^ cells in Lin28a^f/f^ animals (Figure 7F – H). These results indicate that Lin28a in NPCs was necessary for the regulation of adult hippocampal neurogenesis by niche Wnt signaling.

**Figure 7.**
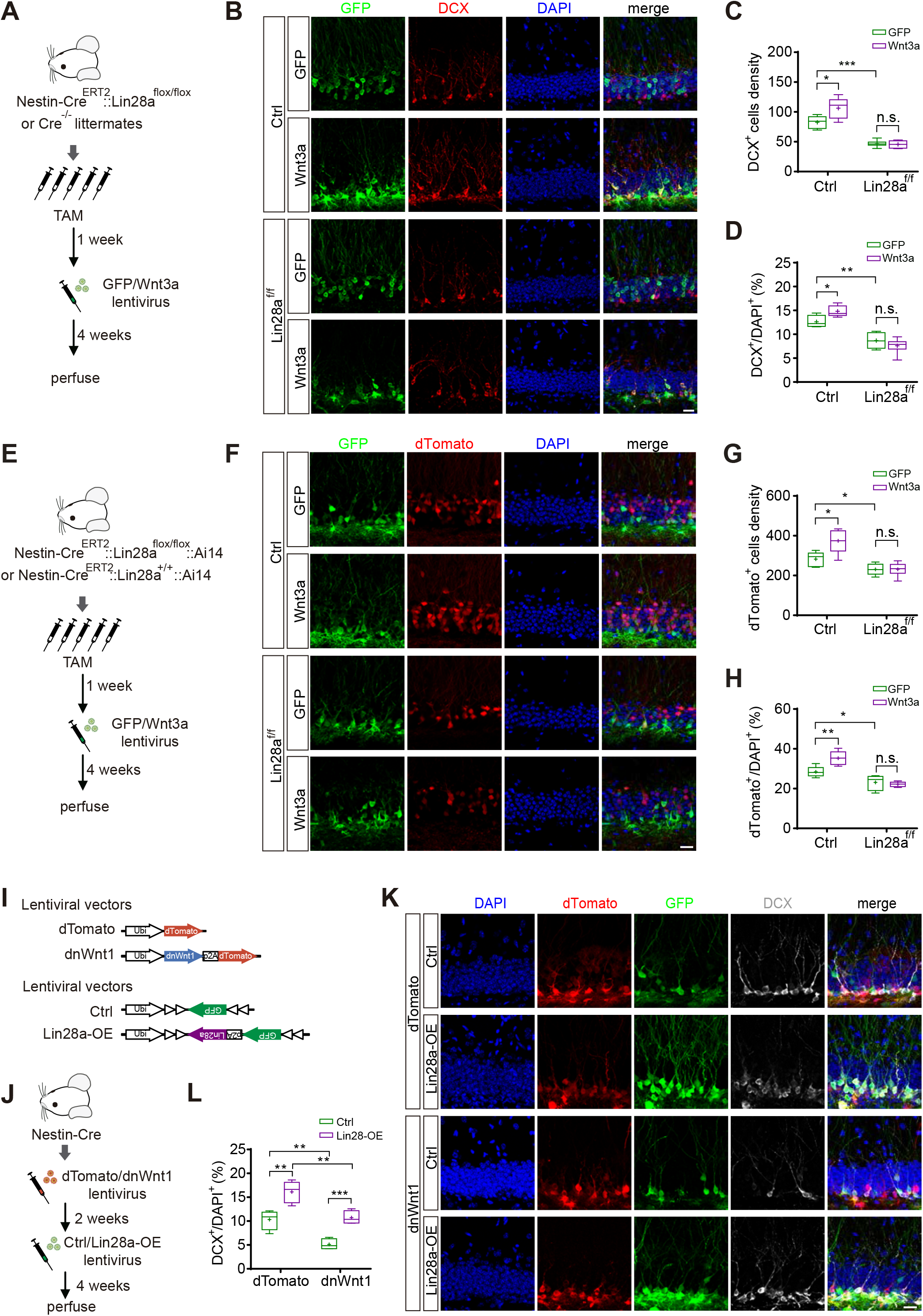
Lin28a is involved in Wnt-mediated regulation of hippocampal neurogenesis. **(A)** Schematics showing that adult Nestin-Cre^ERT2^::Lin28a^flox/flox^ mice (Lin28a^f/f^) or Cre^−/−^ litter mates (Ctrl) were treated with TAM, followed by injection of lentiviral vectors expressing GFP or Wnt3a in the DG, one week later. Four weeks after viral injection, the animals were perfused. **(B)** Confocal images showing DCX^+^ newborn neurons in the DG of Ctrl and Lin28a^f/f^ mice treated with TAM, and injected with GFP or Wnt3a lentiviruses. Scale bar: 20 μm. **(C)** Wnt3a increased the density of DCX^+^ cell in the DG of Ctrl mice, but not in Lin28a^f/f^ mice. (Ctrl: GFP n=4 mice, Wnt3a n=5 mice; two-tailed unpaired t-test, t_7_=2.554, *P=0.0379; Lin28a^f/f^: GFP n=6 mice, Wnt3a n=6 mice; two-tailed unpaired t-test, t10=0.3891, P=0.7054; Ctrl GFP vs. Lin28a^f/f^ GFP: two-tailed unpaired t-test, t_8_=6.341, ***P=0.0002). **(D)** Wnt3a increased the percentage of DCX^+^ cell in the total cells (DAPI^+^) of the GCL in Ctrl mice, but not in Lin28a^f/f^ mice. (Ctrl: GFP n=4 mice, Wnt3a n=5 mice; two-tailed unpaired t-test, t_8_=2.718, *P=0.0299; Lin28a^f/f^: GFP n=6 mice, Wnt3a n=6 mice; two-tailed unpaired t-test, t_10_=1.113, P=0.2918; Ctrl GFP vs. Lin28a^f/f^ GFP: two-tailed unpaired t-test, t_8_=3.877, **P=0.0047). **(E)** Schematics showing that adult Nestin-Cre^ERT2^::Lin28a^flox/flox^::Ai14 mice (Lin28a^f/f^) or Nestin-Cre^ERT2^::Ai14 mice (Ctrl) were treated with TAM, followed by injection of lentiviral vectors expressing GFP or Wnt3a in the DG, one week later. Four weeks after viral injection, the animals were perfused for further analysis. **(F)** Confocal images showing dTomato^+^ adult-born neurons in the DG of Ctrl and Lin28a^f/f^ mice treated with TAM, and injected with GFP or Wnt3a lentiviruses. Scale bar: 20 μm. **(G)** Wnt3a increased the density of dTomato^+^ cell in the DG of Ctrl mice, but not in Lin28a^f/f^ mice. (Ctrl: GFP n=5 mice, Wnt3a n=5 mice; two-tailed unpaired t-test, t_8_=2.921, *P=0.0193; Lin28a^f/f^: GFP n=6 mice, Wnt3a n=6 mice; two-tailed unpaired t-test, t_10_=0.03851, P=0.9700; Ctrl GFP vs. Lin28a^f/f^ GFP: two-tailed unpaired t-test, t_9_=2.705, *P=0.0242). **(H)** Wnt3a increased the percentage of dTomato^+^ cell in the total cells (DAPI^+^) of the GCL in Ctrl mice, but not in Lin28a^f/f^ mice. (Ctrl: GFP n=5 mice, Wnt3a n=5 mice; two-tailed unpaired t-test, t_8_=3.531, **P=0.0077; Lin28a^f/f^: GFP n=6 mice, Wnt3a n=6 mice; two-tailed unpaired t-test, t_10_=0.5937, P=0.5659; Ctrl GFP vs. Lin28a^f/f^ GFP: two-tailed unpaired t-test, t_9_=2.714, *P=0.0238). **(I)** Schematics showing lentiviral vectors expressing dTomato or dnWnt1 (upper panel), and FLEX lentiviral vectors expressing GFP (Ctrl) or GFP-p2A-Lin28a (Lin28a-OE) (lower panel). **(J)** Schematics showing that lentiviral vectors expressing dTomato or dnWnt1 were injected into the DG of adult Nestin-Cre mice, and Cre-dependent Ctrl or Lin28a-OE lentiviral vectors were injected 2 weeks later. Four more weeks later, the animals were perfused. **(K)** Confocal images showing DCX^+^ newborn neurons in the DG of Ctrl and Lin28a^f/f^ mice treated with TAM, and injected with GFP or Wnt3a lentiviruses. Scale bar: 20μm. **(L)** dnWnt1 decreased the percentage of DCX^+^ cell in the total cells (DAPI^+^) of the GCL, whereas upregulating Lin28a in NPCs recovered the number of DCX^+^ cells. (dTomato: Ctrl n=4 mice, Lin28a-OE n=5 mice; two-tailed unpaired t-test, t_7_=3.993, **P=0.0052; dnWnt1: Ctrl n=4 mice, Lin28a-OE n=4 mice; two-tailed unpaired t-test, t_6_=6.199, ***P=0.0008; dTomato Ctrl vs. dnWnt1 Ctrl: two-tailed unpaired t-test, t_6_=4.421, **P=0.0045; dTomato Lin28a-OE vs. dnWnt1 Lin28a-OE: two-tailed unpaired t-test, t_7_=4.181, **P=0.0041).

Previous studies have shown that the deficiency in Wnt signaling results in decreased neurogenesis in the adult hippocampus (Jang et al., 2013; Seib et al., 2013). To test whether the overexpression of Lin28a is sufficient to recover impaired neurogenesis in Wnt-deficient hippocampus, we constructed a lentiviral vector expressing a dominant negative Wnt1 (dnWnt1) (Figure 7I), which has been shown to inhibit Wnt signaling and decrease neurogenesis (Choi et al., 2018; Lie et al., 2005). We also constructed another lentiviral vector that contains a FLEX GFP-p2A-Lin28a, so that we could overexpress Lin28a in the presence of Cre recombinase (Figure 7I). We injected the lentivirus expressing dnWnt1-dTomato or dTomato only into the DG of Nestin-Cre mice, and infused lentivirus expressing FLEX GFP-p2A-Lin28a two weeks later to overexpress Lin28a in the Nestin^+^ NPCs (using lentivirus expressing FLEX-GFP as Ctrl). Four more weeks later, we perfused the mice and quantified the number of DCX^+^ cells in the DG of these animals (Figure 7J). We found expression of dnWnt1 indeed decreased the number of DCX^+^ newborn neurons in the DG, whereas upregulation of Lin28a in NPCs increased the number of DCX^+^ cells in dTomato group and recovered the number of DCX^+^ cells in the dnWnt1 group (Figure 7K and L). These data suggest that overexpressing Lin28a in NPCs was sufficient to restore neurogenesis in Wnt-deficient hippocampal neurogenic niche.

### Overexpression of Lin28a in NPCs restores neurogenesis in the aging hippocampus and enhances pattern separation

Since the overexpression of Lin28a could upregulate the proliferation of NPCs and facilitate the integration of newborn cells, we wondered whether overexpression of Lin28a in NPCs could enhance pattern separation by increasing hippocampal neurogenesis. We injected Cre-dependent lentivirus expressing FLEX GFP-p2A-Lin28a (Lin28a-OE) into the DG of young adult Nestin-Cre mice (2mo) to overexpress Lin28a in a large population of existing NPCs in SGZ, while virus expressing GFP was use as control (Ctrl) (Figure 8A and B). Six weeks later, the mice showed significantly increased DCX^+^ newborn neurons when Lin28a is specifically overexpressed in Nestin^+^ NPCs and their progenies, compared to Ctrl mice (Figure 8C and D). Lin28a-OE mice started to show discrimination between contexts A and B in training session 2, while Ctrl mice started to show discrimination in session 4 (Figure 8E), suggesting Lin28a-OE mice exhibited enhanced ability of pattern separation.

**Figure 8.**
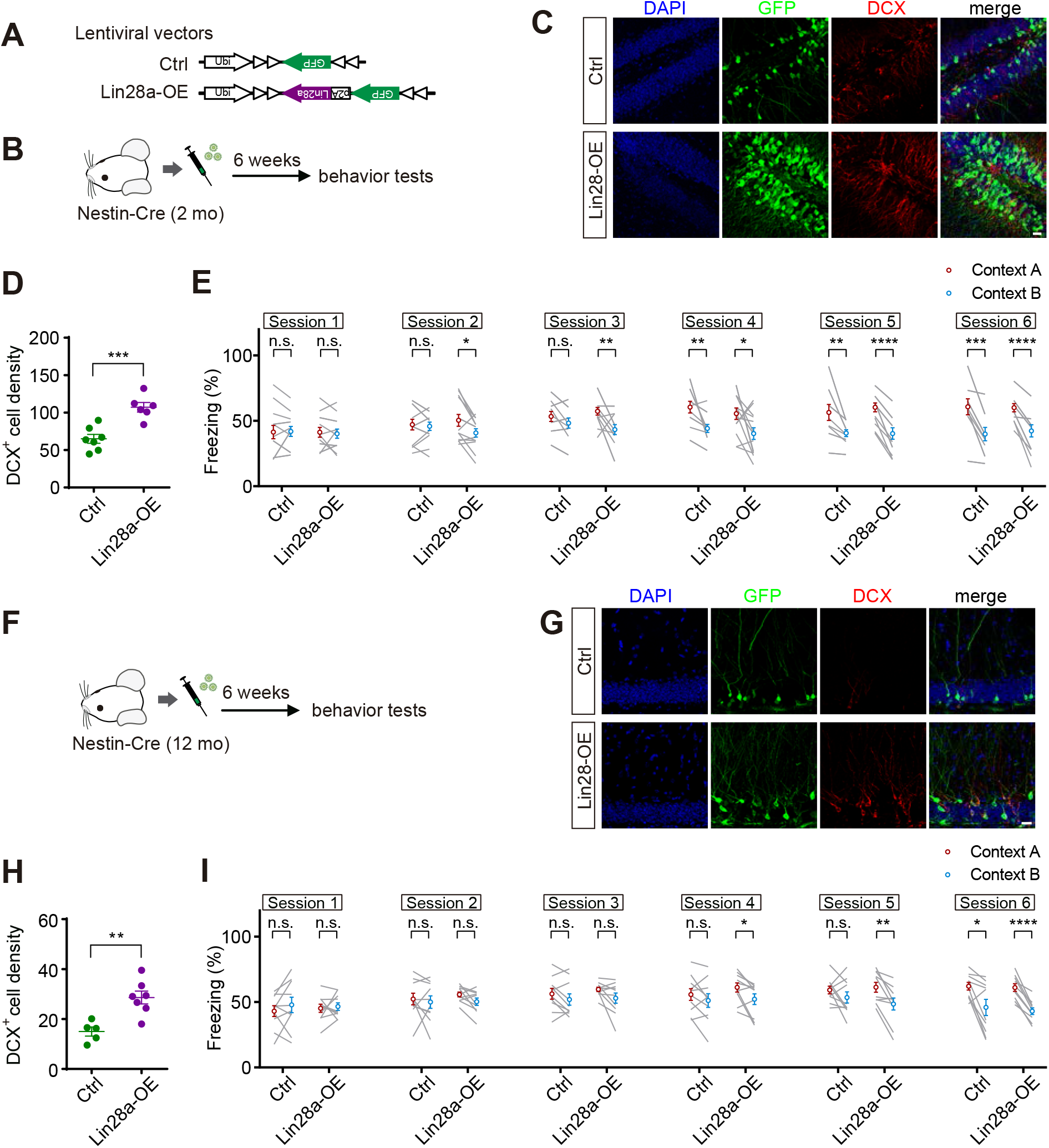
Overexpression of Lin28a in NPCs restores neurogenesis in the aging hippocampus and enhances pattern separation. **(A)** Diagram showing FLEX GFP (Ctrl) or GFP-p2A-Lin28a (Lin28a-OE) lentiviral vectors. **(B)** The lentiviruses were injected into the DG of young adult Nestin-Cre mice (2 mo), followed by behavioral tests 6 weeks later. **(C)** Confocal images showing DCX^+^ cells in the DG of Nestin-Cre mice injected with FLEX lentivirus expressing GFP (Ctrl) or GFP-p2A-Lin28a (Lin28a-OE). Scale bar: 20 μm. **(D)** Density of DCX^+^ cells in the DG of Ctrl and Lin28a-OE mice. (Ctrl n=7 mice, Lin28a-OE n=6 mice; two-tailed unpaired t-test, t_11_=4.807, ***P=0.0005). **(E)** The freezing of Ctrl and Lin28a-OE mice in contexts A and B in test sessions 1 through 6. (Ctrl mice n=13, Lin28a-OE mice n=13; two-tailed paired t-test; session 1: Ctrl t_12_=0.1723, P=0.8661; Lin28a-OE t_12_=0.4251, P=0.6783; session 2: Ctrl t_12_=0.5160, P=0.6152; Lin28a-OE t_12_=2.922, *P=0.0128; session 3: Ctrl t_12_=2.033, P=0.0648; Lin28a-OE t_12_=3.121, **P=0.0088; session 4: Ctrl t_12_=3.662, **P=0.0033; Lin28a-OE t_12_=2.470, *P=0.0295; session 5: Ctrl t_12_=3.925, **P=0.0020; Lin28a-OE t_12_=8.950, ****P<0.0001; session 6: Ctrl t_12_=5.391, ***P=0.0002; Lin28a-OE t_12_=6.209, ****P<0.0001). **(F)** Diagram showing Cre-dependent Ctrl or Lin28a-OE lentiviruses were injected into the DG of aging Nestin-Cre mice (12 mo), followed by behavioral tests six weeks later. **(G)** Confocal images showing DCX^+^ cells in the DG of aging Nestin-Cre mice (12 mo) 6 weeks after the injection of FLEX lentivirus expressing GFP (Ctrl) or GFP-p2A-Lin28a (Lin28a-OE). Scale bar: 20 μm. **(H)** Density of DCX^+^ cells in the DG of Ctrl and Lin28a-OE mice. (Ctrl n=5 mice, Lin28a-OE n=7 mice; two-tailed unpaired t-test, t_10_=3.970, **P=0.0026). **(I)** The freezing of Ctrl and Lin28a-OE mice in contexts A and B in test sessions 1 through 6. (Ctrl mice n=10, Lin28a-OE mice n=10; two-tailed paired t-test; session 1: Ctrl t_9_=1.052, P=0.3203; Lin28a-OE t_9_=0.3730, P=0.7178; session 2: Ctrl t_9_=0.4910, P=0.6352; Lin28a-OE t_9_=2.131, P=0.0620; session 3: Ctrl t_9_=1.606, P=0.1426; Lin28a-OE t_9_=1.979, P=0.0792; session 4: Ctrl t_9_=1.335, P=0.2146; Lin28a-OE t_9_=2.716, *P=0.0237; session 5: Ctrl t_9_=1.292, P=0.2287; Lin28a-OE t_9_=3.840, **P=0.0040; session 6: Ctrl t_9_=3.214, *P=0.0106; Lin28a-OE t_9_=8.339, ****P<0.0001).

To test whether overexpressing Lin28a could enhance neurogenesis in aging animal and improve their ability of pattern separation, we injected the above-mentioned lentiviruses into the DG of aging Nestin-Cre mice (12 mo) (Figure 8F). Six weeks later, we found that overexpression of Lin28a in the NPCs in aging mice showed increased number of DCX^+^ newborn neurons in the DG, compared to Ctrl mice (Figure 8G and H). Consistent with the data shown in Figure 2B, the aging mice in the Ctrl group exhibited poor contextual discrimination, as the freezing time of these animals in contexts A and B did not show difference till session 6 (Figure 8I). Interestingly, when Lin28a was overexpressed in the remaining Nestin^+^ NPCs in the DG of aging mice, the animals started to discriminate contexts A and B in session 4, much sooner than the Ctrl group, suggesting a restored ability of pattern separation (Figure 8I).

These results demonstrate that increasing the expression of Lin28a in NPCs promoted neurogenesis, thus leading to enhanced pattern separation. Moreover, overexpressing Lin28a in hippocampal NPCs in aging mice could rescue their weakened ability of pattern separation by increasing neurogenesis in the DG.

## Discussion

### Lin28a regulates neurogenesis in the adult hippocampus

RNA binding protein Lin28 has been shown to play important roles in a variety of biological processes, through let-7-dependent and independent pathways (Shyh-Chang and Daley, 2013). During the early development of the nervous system, Lin28 is highly expressed in neural stem/progenitor cells, whereas its expression level drops sharply as neuronal differentiation proceeds (Yang et al., 2015). Interestingly, our results showed that Lin28a remained existent in NPCs and DGCs in the adult hippocampus, suggesting its potential biological role in these cells.

Loss of Lin28a in hippocampal NPCs decreased neurogenesis, whereas the overexpression of Lin28a increased neurogenesis, by enhancing the proliferation of NPCs. This result is in agreement with previous study showing that, Lin28a knockout in the developing brain results in microcephaly whereas overexpressing Lin28a in Nestin^+^ NPCs leads to enlarged brain size (Yang et al., 2015). Our results also showed that, the overexpression of Lin28a in NPCs enhanced neurogenesis mainly by increasing the number of cell cycles, but not by altering the survival of newborn cells. Further analysis showed that overexpression of Lin28a decreased the proportion of newborn cell differentiated towards astrocytes, in consistence with a previous report showing that Lin28 suppresses glial fate of NPCs *in vitro* (Balzer et al., 2010).

In addition, we found Lin28a was not only expressed in NPCs, but also in DG neurons. Loss of Lin28a in newborn neurons inhibited, while the overexpression of Lin28a facilitated, the development and functional integration of newborn neurons, suggesting Lin28a not only regulates the proliferation of NPCs, but also controls the growth of newborn neurons in the adult hippocampus.

However, opposite to our results, a recent study showed that overexpression of Lin28 in the SVZ NPCs decreased the number of new neurons in the olfactory bulb of postnatal mice (Romer-Seibert et al., 2019). This result may be attributed to that the overexpression of Lin28 repressed let-7, which is important for the migration of neuroblasts (Petri et al., 2017), thus decreasing the number of new neurons arriving in the olfactory bulb. Although it is reported that short-distance lateral dispersion is required for the integration of newborn neurons in the DG (Wang et al., 2019), we did not observe developmental or integrating defects in newborn neurons when Lin28a is overexpressed.

It has been shown that Lin28a and Lin28b are both required and play overlapping functions in regulating NPC proliferation and brain development during early development (Yang et al., 2015). We did not detect Lin28b expression in the adult DG using RNAscope, either in Nestin^+^ NPCs or in Prox1^+^ DGCs, suggesting differential expression of Lin28a and Lin28b in the adult brain. It is reported that Lin28b knockout mice exhibit no detectable phenotypes (Shinoda et al., 2013), whereas loss of Lin28a results in reduced brain size at birth (Yang et al., 2015). These suggest that Lin28a serves as the dominant Lin28 homolog regulating neurogenesis in both developing and adult brains. Moreover, it has been reported that the expression of Lin28a, but not Lin28b, is regulated by Wnt-β-catenin signaling (Cai et al., 2013), suggesting Lin28a may play more important roles rather than Lin28b in regulating neurogenesis in the presence of extracellular Wnt signals.

### Decrease of Lin28a expression is involved in aging-associated decline of neurogenesis

Hippocampal neurogenesis decreases in mammals with aging (Kuhn et al., 1996; Spalding et al., 2013), associated with and contributing to cognitive declines in aging (Lee et al., 2012; Seib et al., 2013). Multiple pathways have been revealed to contribute to age-related decline of adult neurogenesis (Meyers et al., 2016; Miranda et al., 2012; Seib et al., 2013; Yang et al., 2017; Yousef et al., 2015). Wnt signaling pathway has been shown to be of essential importance for the regulation of multiple stages of adult hippocampal neurogenesis (Lie et al., 2005; Schafer et al., 2015). Both existing in the neurogenic niche and changing with aging or activity, Wnt ligands promote (Lie et al., 2005; Song et al., 2002), while Wnt antagonists inhibit neurogenesis (Jang et al., 2013; Seib et al., 2013), thus contributing to aging-associated decline of neurogenesis.

Interestingly, we observed a decreased expression of Lin28a in the hippocampal NPCs with aging, coincident with the increased Wnt antagonist Dkk1 in the niche (Seib et al., 2013). Furthermore, we found the upregulation of niche Wnt3a or overexpression of Ctnnb1 in the NPCs increased the expression of Lin28a in hippocampal NPCs, in consistence with a previous study that Wnt-β-catenin signaling regulates the expression of Lin28a, by direct binding of β-catenin to Lin28a promoter (Cai et al., 2013). A recent study also showed that Wnt-β-catenin signaling regulates the proliferation and neurogenic potential of retinal müller glia cells via Lin28/let-7 pathway (Yao et al., 2016). These clues suggest that Lin28a serves as a downstream mechanism underlying the regulation of neurogenesis by niche Wnt signals.

In the adult hippocampus, our results showed that Lin28a regulated the proliferation of NPCs and the development of new neurons, consistent with the regulation of neurogenesis by Wnt signals (Jang et al., 2013; Lie et al., 2005; Seib et al., 2013). Especially, loss of Lin28a from NPCs reduced neurogenesis and inhibited the development of new neurons, mimicking the effects of increased Wnt antagonists in the neurogenic niche (Jang et al., 2013; Seib et al., 2013). Thus, in the aging brain, with the increasing niche Wnt antagonists (Seib et al., 2013), decreased Lin28a expression in NPCs contributes to the declining neurogenesis.

Importantly, lacking of Lin28a in NPCs blocked the enhanced neurogenesis by Wnt3a, suggesting that Lin28a is essential for NPCs to respond to local Wnt signals in the adult hippocampal neurogenic niche. On the other hand, upregulation of Wnt antagonist dnWnt1 in the neurogenic niche, which has been shown to block Wnt signaling (Choi et al., 2018; Lie et al., 2005), significantly decreased adult neurogenesis in the DG; whereas the overexpression of Lin28a in NPCs could rescue neurogenesis in this Wnt-deficient microenvironment. Consistently, in aging animals, the overexpression of Lin28a in the remaining NPCs could at least partially restore hippocampal neurogenesis.

It has been shown that young adult-born neurons in the DG are essential for pattern separation (Nakashiba et al., 2012; Sahay et al., 2011a). In human, pattern separation undergoes an age-related decline in performance, correlating with the decay of neurogenesis in the DG (Stark et al., 2010). Knocking out Lin28a from NPCs resulted in decreased neurogenesis and impaired pattern separation, recapitulating the phenotype in aging animals. Overexpression of Lin28a in NPCs could increase neurogenesis, and thus enhance pattern separation in both young and aging mice. These results suggest that pattern separation is susceptible to the changes in hippocampal neurogenesis, which is regulated by Lin28a level in hippocampal NPCs.

Taken together, our study revealed that Lin28a regulates neurogenesis in response to Wnt signals in the adult hippocampus *in vivo*, suggesting that Lin28a acts as one important link between niche Wnt ligands/antagonist and neurogenesis in the aging brain. As other target genes of Wnt-β-catenin signaling through TCF/LEF, such as *Neurod1* and *Prox1*, have also been shown to regulate adult neurogenesis (Gao et al., 2009; Kuwabara et al., 2009; Lavado et al., 2010; Lie et al., 2005; Miranda et al., 2012), further studies will be needed for understanding possible interactions between these molecules in NPCs.

## Materials and Methods

### Animals

All procedures were approved by the Animal Care and Use Committees at the Zhejiang University School of Medicine, and conducted in accordance with the policies of institutional guidelines on the care and use of laboratory animals.

All mice were bred in the animal facility at the Zhejiang University School of Medicine, and maintained at a 12 hr light/dark cycle. Mice were group housed (3–4 per cage) in transparent plastic cages (35 × 15 × 20 cm), with free access to normal food and water unless otherwise specified. Males and females were equally used for experiments.

Lin28a^flox/flox^ mice (Stock No. 023913), Nestin-Cre mice (Stock No. 003771), Nestin-Cre^ERT2^ mice (Stock No. 016261) and Ai14 reporter line (Stock No. 007914) were from Jackson Labs; Nestin-GFP mice (Stock No. RBRC06355) were from RIKEN.

### Tamoxifen administration

Tamoxifen (TAM) (Sigma, T5648) was dissolved in vegetable oil containing 0.1% absolute ethanol to a concentration of 100mg/ml.

For conditional knocking out Lin28a from Nestin^+^ NPCs in Nestin-Cre^ERT2^::Lin28a^flox/flox^ mice, or labeling and knocking out Lin28a from Nestin^+^ NPCs in Nestin-Cre^ERT2^::Lin28a^flox/flox^::Ai14 mice, TAM was administered i.p. at the dose of 160mg/kg every three days for five times.

### Construction of viral vectors

#### 1) Lin28a retroviral and lentiviral vectors

Mouse Lin28a mRNA CDS sequence was obtained from GenBank (NCBI Reference Sequence: NM_145833.1, CCDS18761.1), synthesized and inserted into a pUX-GFP retroviral plasmid vector to make pUX-GFP-p2A-Lin28a. pUX-GFP was used as control vector.

GFP-p2A-Lin28a fragment was then subcloned and inserted in a reversed direction into a pUX-FLEX retroviral plasmid vector to make a Cre-dependent retroviral plasmid vector pUX-FLEX-GFP-p2A-Lin28a. pUX-FLEX-GFP was used as control vector.

GFP-p2A-Lin28a fragment was then subcloned and inserted in a reversed direction into a pHage-FLEX lentiviral plasmid vector to make a Cre-dependent lentiviral plasmid vector pHage-FLEX-GFP-p2A-Lin28a. pHage-FLEX-GFP was used as control vector.

#### 2) Truncated Lin28a (tLin28a) and mutant Lin28a (mutLin28a) retroviral vectors

A truncated Lin28a sequence (1 to 408 of Lin28a cDNA sequence) was subcloned into pUX-GFP plasmid vector to make pUX-GFP-p2A-tLin28a. GFP-p2A-tLin28a fragment was then cut and inserted in a reversed direction into a pUX-FLEX retroviral plasmid vector to make pUX-FLEX-GFP-p2A-tLin28a.

Lin28a DNA sequence with two point mutations (to change H147 and H169 to alanines in the protein) was synthesized and subcloned into pUX-GFP plasmid vector to make pUX-GFP-p2A-mutLin28a. GFP-p2A-mutLin28a fragment was then cut and inserted in a reversed direction into a pUX-FLEX retroviral plasmid vector to make pUX-FLEX-GFP-p2A-mutLin28a.

#### 3) Wnt3a lentiviral vector

Mouse Wnt3a mRNA CDS sequence was obtained from GenBank (NCBI Reference Sequence: NM_009522.2, CCDS 24766.1), synthesized and then subcloned and inserted in a reversed direction into a pHage-GFP lentiviral plasmid vector to make a lentiviral plasmid vector pHage-Wnt3a-p2A-GFP.

#### 4) dnWnt1 lentiviral vector

A dominant negative mouse Wnt1 (dnWnt1) sequence, as previously described (Hoppler et al., 1996), was synthesized and then subcloned and inserted in a reversed direction into a pHage-GFP lentiviral plasmid vector to make a lentiviral plasmid vector pHage-dnWnt1-p2A-dTomato.

#### 5) Ctnnb1 lentiviral vector

Mouse Ctnnb1 mRNA CDS sequence was obtained from GenBank (NCBI Reference Sequence: NM_007614.3, CCDS23630.1), synthesized and then subcloned and inserted in a reversed direction into a pHage-FLEX-GFP lentiviral plasmid vector to make a lentiviral plasmid vector pHage-FLEX-Ctnnb1-p2A-GFP.

#### 6) m/c/nXFP retroviral vectors

Membrane RFP (mRFP) and nucleus CFP (nCFP) sequences were inserted in a reversed direction into pUX-FLEX retroviral vector to make the pUX-FLEX-mRFP and pUX-FLEX-nCFP vectors, respectively. Above mentioned pUX-FLEX-GFP vector was used as pUX-FLEX-cGFP (cytosol GFP).

### Virus production

As previously described (Gu et al., 2012), each retroviral or lentiviral backbone plasmid was co-transfected to 293T cells with helper plasmids for retrovirus or lentivirus using Lipofectamine 2000 (Invitrogen). Culture medium was collected, then virus was purified and concentrated by ultracentrifugation.

### Virus injection

Stereotactic viral injections were performed in accordance with the Guidelines by Zhejiang University Animal Care and Use Committees. As previously described (Gu et al., 2012), mice were anesthetized using isoflurane and mounted on a stereotaxic machine. Viral particles were then injected using a 1 μl Hamilton syringe bilaterally into the DG (coordinates: 2.0 mm caudal to bregma, 1.6 mm lateral from the midline and 2.5 mm ventral; 3.0 mm caudal to bregma, 2.6 mm lateral from the midline and 3.2 mm ventral). Mice were then returned to their home cages after waking up, and then housed under standard conditions.

### Immunostaining, confocal imaging and image analysis

As previously described (Gu et al., 2012), mice were deeply anesthetized and perfused transcardially with PBS and then 4% PFA. Brains were removed, fixed overnight in PFA and then transferred to a 30% (w/v) sucrose solution and stored at 4 °C. Brains were sectioned into 50-μm coronal sections covering the full anterior-posterior extent of the hippocampus. Regular immunohistochemistry was performed using primary antibodies to Lin28a (Cell Signaling Technology, 8641, 1:50), Lin28 (Abcam, ab63740, 1:50), Prox1 (Abcam, ab199359, 1:500), NeuN (Abcam, ab104224, 1:1000), Iba-1(Wako, 019-19741, 1:1000), GFAP(Abcam, ab4674, 1:1000), s100β (Abcam, ab52642, 1:1000), MCM2 (Cell Signaling Technology, 4007, 1:500), DCX (Cell Signaling Technology, 4604, 1:1000) or GFP (Abcam, ab12218, 1:3000), overnight at 4 °C, followed by incubation with secondary antibodies (Cy2-conjugated donkey antibody to chicken (Jackson Immune, 703-225-155, 1:1000), Cy3-conjugated donkey antibody to rabbit (Jackson Immune, 711-165-152, 1:1000), Cy5-conjugated donkey antibody to rabbit (Jackson Immune, 711-175-152, 1:1000) or Cy5-conjugated donkey antibody to chicken (Jackson Immune, 703-175-155, 1:1000) for 2 hours at room temperature (25°C), in the presence of 2% donkey serum, 1% bovine serum albumin and 0.2% (w/v) Triton X-100. Sections were mounted on slides with Fluoromount-G anti-fade medium containing DAPI (SouthernBiotech). Images of z series stacks were taken on an Olympus FV3000 confocal microscope. Images were analyzed using Image J or Imaris software.

For analyzing the neurogenesis in the DG, the density and percentage of MCM2^+^, DCX^+^ or dTomato-labeled cells were calculated and analyzed. For calculating the density of cells, we flattened the SGZ and projected the MCM2^+^, DCX^+^ or dTomato^+^ cell body to the plane of the flattened SGZ, and the density of each type of cell was calculated as number of MCM2^+^, DCX^+^ cells per 100×100 μm^2^ area of the flattened SGZ. The percentage of MCM2^+^, DCX^+^ or dTomato^+^ cells in the dentate gyrus was calculated as the proportion of each types of cells in the total number of cells (DAPI^+^) in the cell body layer of the analyzed DG region. The cell counting covered the whole dentate gyrus in each section, and sections from dorsal through ventral part of the hippocampus were imaged, analyzed, and then averaged for each animal.

### Clonal labeling and analysis

Engineered self-inactivating murine oncoretroviruses were used to deliver genes of interest specifically to proliferating cells and their progeny (Ge et al., 2006; van Praag et al., 2002). To label clusters of newborn cells generated from individual NPCs, we injected FLEX retrovirus expressing GFP or overexpressing Lin28a into the DG of Nestin-Cre mice, thus resulting in sparse labeling of progenies of individual dividing Nestin^+^ neural stem cells, as we previous described (Kirschen et al., 2017). Brains were sliced into 60 μm sections and GFP was enhanced by staining. Sections were then mounted on slides with Fluoromount-G anti-fade medium with DAPI. For analysis, we only selected isolated cell clusters, and cells within 100 µm distance were considered belonging to the same cell cluster.

To avoid mixture of adjacent cell clusters, we used a combination of FLEX m/n/cXFP retroviral vectors to label different cell clusters with different combination of colors. m/n/cXFP retroviruses were mixed and injected into the DG of Nestin-Cre mice, and the animals were perfused 7 days after viral injection. Brains were sliced into 60 μm sections, which were then mounted on slides with Fluoromount-G anti-fade medium without DAPI (SouthernBiotech). For analysis, only adjacent labeled cells with the same combination of colors were considered belonging to the same cell cluster. For Lin28a, tLin28a or mutLin28a groups, we mixed pUX-FLEX-GFP-p2A-Lin28a, pUX-FLEX-GFP-p2A-tLin28a or pUX-FLEX-GFP-p2A-mutLin28a with pUX-FLEX-mRFP and pUX-FLEX-nCFP retroviruses for combinations. Only cells expressing GFP were considered as overexpressing Lin28a, tLin28a or mutLin28a.

### Labeling the division of NPCs

Cre-dependent retrovirus (pUX-FLEX-GFP or pUX-FLEX-GFP-p2A-Lin28a) was injected into the DG of adult Nestin-Cre mice. 48 hours after viral injection, BrdU (Sigma, B5002) was administered (i.p., 100mg/kg) every 6 hours for 5 times. Seven days after viral injection, mice were perfused and brain sections were stained using primary antibodies to GFP (Abcam, ab12218, 1:3000), BrdU (Abcam, Ab6326, 1:1000) and MCM2 (Cell Signaling Technology, 4007, 1:500) over night at 4 °C, followed by incubation with secondary antibodies. Images of GFP-labeled cells were taken on an Olympus FV3000 confocal microscope, and were analyzed for colocalization of GFP, BrdU and MCM2, using Image J or Imaris.

### Labeling apoptotic newborn cells

Retrovirus expressing GFP (Ctrl) or GFP-p2A-Lin28a (Lin28a-OE) was injected into the DG of adult C57 mice. 3 days after viral injection, mice were perfused and brain sections were stained using primary antibodies to GFP (Abcam, ab12218, 1:3000) and aCas-3 (Abcam, ab13847, 1:200) over night at 4 °C, followed by incubation with secondary antibodies. Images of GFP-labeled cells were taken on an Olympus FV3000 confocal microscope, and colocalization of GFP and aCas-3 were analyzed for using Image J or Imaris.

### RNAscope in situ hybridization

Brain slices (14μm) were prepared as described above. In situ hybridization was performed using RNAscope multiplex fluorescent reagent kit (Advanced Cell Diagnostics) according to manufacturer’s standard instructions. Ethanol was used for gradient dehydration of the sections. Probes were designed by ACDbio (probes : Mm-Lin28a, 437121; Mm-Sox2-C3, 401041-C3; Neg-ctrl, 320871; Pos-ctrl, 320881).

Additional immunostaining was done following the detection of mRNA with standard protocols as described above. Images were obtained using an Olympus FV3000 confocal microscope. For the quantification of mRNA expression level in single cells, the number of dots in each cell of interest were counted using ImageJ software. To minimized the influence of non-specific signals, RNAscope using the negative control probe (Neg-ctrl) was also performed at the same time on the same sets of samples, and the average number of dots in each cell obtained using Neg-ctrl probes was subtracted from the number of dots obtained using target probes.

### Western blot analysis

To detect the Lin28a and other protein levels in the DG, mice were anesthetized with isoflurane (1.5-3.0%) and the brain was removed and cut into acute slices (300μm) in cold PBS. The DG was dissected and collected, then homogenized and centrifuged at 12000 r.p.m. at 4℃ for 10minutes. Each sample contained DG tissues from 3 animals. Protein concentration was detected by the BCA Protein Assay Kit (Beyotime, P0011). Samples were diluted in 5× loading buffer and were boiled at 100℃ for 10min. Proteins were then separated in 12% gel by SDS-PAGE and transferred to PVDF membranes (Merck Millipore, ISEQ00010), which were then block with 5% milk in TBS-T buffer at room temperature for 1 hour. Membranes were incubated with primary antibodies to Lin28a (Cell Signaling Technology, 8641, 1:500), rabbit-anti-Lin28 (Abcam, ab63740, 1:500) or rabbit-β-catenin (Cell Signaling, 8480, 1:1000) at 4°C overnight. Membranes were then incubated with horseradish peroxidase-conjugated goat-anti-rabbit-IgG (Earthox, 1:5000) at room temperature for 1 hour. Bands were visualized with the ECL kit (Thermo, 34077) and imaged using the GE (AI600). β-tubulin was used as internal control. Quantitative analysis was performed with ImageJ software. All experiments were repeated for at least three times.

### Whole cell electrophysiological recordings

Acute brain sections were made 3 weeks after retroviral injection, and electrophysiological recordings were performed at 32–34°C, as previously described (Ge et al., 2006). Briefly, mice were deeply anesthetized with isoflurane, and brain was removed and acute brain slices were prepared. GFP-labeled neurons were identified under fluorescent microscope and whole-cell patchclamp recordings were performed. Miniature excitatory postsynaptic currents (mEPSCs) were recorded in the presence of tetrodotoxin (1μM) and picrotoxin (100 μM) at a holding potential of −70mV. Miniature events were automatically detected and then analyzed using Clampfit software.

### Behavior procedures

#### 1) Contextual fear conditioning (CFC) and test

Mice were trained in a conditioning chamber (18 cm×18 cm×30 cm; Coulbourn, Whitehall, PA), containing a stainless-steel shock-grid floor (context A). During the training, mice were placed in the chamber and received a single foot shock (0.75 mA, 2 s duration) after 2 min. Mice were taken out of the training chamber 1 min after the foot shock and placed back into their home cages. After training, mice were housed under standard conditions with 12 hr light/dark cycle till test. 24 hours later, mice were placed back in context A for 5 min, without foot shock. Freezing behavior of animals was monitored via overhead cameras and measured by an automated scoring system.

#### 2) Pattern separation trainings and tests

Mice received CFC training in context A on day 0. From day 1, mice were exposed to both context A and context B, 4 hours apart with random order, for 12 consecutive days. In context A, mice were exposed to the context for 3-min followed by a 2-s foot shock (0.75 mA) and then 1-min stay. In context B, mice were allowed to stay for 3 min without foot shock. Context B is a similar context sharing many features with context A, including the same stainless-steel grid floor and roof, but only different with striped pattern in front and back walls. The freezing of mice in two consecutive days with contexts A and B in different orders was averaged as one session to minimize variation.

#### 3) Novel location and novel object recognition tests

For novel location recognition test, mice were placed in a normally illuminated box (45 cm × 45 cm × 35 cm) with an overhead camera, and allowed for habituation for 10 min and then put back to homecage. One hour later, two identical objects were place close to two corners in the box; mice were put in the box and allowed to freely explore for 10 min. Another hour later, before the test session, one of the two identical objects was place at a different location (novel location), while the other object was not moved; mice were put back in the box for 10 min, and the time of their exploration on the two objects were recorded and analyzed.

For novel object recognition test, mice were placed in a normally illuminated box (45 cm × 45 cm × 35 cm) with an overhead camera, and allowed for habituation for 10 min and then put back to homecage. One hour later, two identical objects were place close to two corners in the box; mice were put in the box and allowed to freely explore for 10 min. Another hour later, before the test session, one of the two identical objects was replace with a different object (novel object), while the other object was not changed or moved; mice were put back in the box for 10 min, and the time of their exploration on the two objects were recorded and analyzed.

The discrimination index was calculated as: (Time_novel_ – Time_familiar_)/(Time_novel_ + Time_familiar_).

### Statistical analysis

Data were analyzed using GraphPad Prism 8.0 software. Statistical analysis was carried out using two-tailed unpaired t-tests or otherwise indicated in the figure legends. For analyzing size of cell clusters, two-tailed unpaired t-tests were performed for the average clonal size, following Kolmogorov-Smirnov tests for the cumulative distributions. For the statistical analysis of the discrimination index in the pattern separation tests, two-way RM ANOVA were used to analyze the difference between two groups. Data were presented using box-and-whisker plots or scatter plots. For box-and-whisker plots, whiskers represent the minimum and maximum, the box represent the upper and lower quartiles, the line in the middle represents the median, and the “+” represents the mean. For the scatter plots, data are presented as the mean ± SEM. Statistical significance was considered when P<0.05.

## Acknowledgments

This work was supported by the National Key R&D Program of China (2017YFA0104200), the Zhejiang Provincial Natural Science Foundation of China (LR17C090001), the National Natural Science Foundation of China (32071021) and startup funds to Y.G.; and the National Natural Science Foundation of China (32070975), the Zhejiang Provincial Natural Science Foundation of China (R21C090002) to La. W. We are grateful to the Core Facilities of Zhejiang University School of Medicine for technical assistance.

## Author contributions

Z.H., La.W. and Y.G. designed all the experiments; Z.H. conducted most of the experiments, including most plasmid construction, virus production, viral injection, imaging, electrophysiology, behavioral experiments and related analysis; J.M. did western blots and analysis; H.Y. constructed XFP plasmid vectors and some imaging; X.L. and C.W. did some imaging and analysis; Li.W. provided Ai14 mouse line; B.S. provided Nestin-Cre and Nestin-GFP mouse lines; Z.C. provided Nestin-Cre^ERT2^ mouse line; Z.H., J.M., La.W. and Y.G. discussed the results and wrote the manuscript. All authors read and discussed about the manuscript.

## Expanded View Figure legends

**Figure EV1. Expression of Lin28a in the dentate gyrus. (A)** Confocal image showing Lin28a protein exists in the DG of adult Nestin-GFP mice. Scale bar: 100 μm. **(B)** Confocal images showing Lin28a protein exists in the cells locate in the GCL and SGZ of wildtype (WT) mice, but is absent from the GCL and SGZ of Nestin-Cre::Lin28a^flox/flox^ (Lin28a^f/f^) mice. Scale bar: 10 μm.

**Figure EV2. Aging resulted in decreased hippocampal neurogenesis, and the effects on animals’ behaviors. (A)** Experimental paradigm for novel location recognition test. **(B)** The performance of 2 mo and 10 mo mice in the novel location recognition test. (2 mo n=10 mice, two-tailed paired t-test, t_9_=4.458, **P=0.0016; 10 mo n=10 mice, two-tailed paired t-test, t_9_=0.9220, P=0.3806). **(C)** 10 mo mice showed impaired discrimination between novel and familiar locations. (2 mo n=10 mice, 10 mo n=10 mice, two-tailed unpaired t-test, t_18_=3.193, **P=0.0050). **(D)** Experimental paradigm for novel object recognition test. **(E)** 10 mo mice did not show impaired novel object recognition. (2 mo n=10 mice, two-tailed paired t-test, t_9_=6.184, ***P=0.0002; 10 mo n=10 mice, two-tailed paired t-test, t_9_=4.439, **P=0.0016). **(F)** 2 mo and 10 mo mice showed similar discrimination in novel object test. (2 mo n=10 mice, 10 mo n=10 mice, two-tailed unpaired t-test, t_18_=1.037, P=0.3134). **(G)** Experimental paradigm for contextual fear conditioning and test. Mice were trained in context A and received a footshock, and freezing was tested 24 hours later in the same context. **(H)** 2 mo and 10 mo mice showed similar freezing in single-trial contextual fear memory test. (2 mo n=10 mice, 10 mo n=10 mice, two-tailed unpaired t-test, t_18_=0.3124, P=0.7584). **(I)** Representative confocal images showing DCX^+^ cells in the DG of 2 mo and 10 mo mice. Scale bar: 20 μm. **(J)** Density of DCX^+^ cells in the DG of 2 mo and 10 mo mice. (2 mo n=4 mice, 10 mo n=4 mice, two-tailed unpaired t-test, t_6_=5.044, **P=0.0.0023).

**Figure EV3. Loss of Lin28a from NPCs resulted in decreased neurogenesis, and the effect on animals’ behaviors. (A)** Experimental paradigm for contextual fear conditioning and test. Six weeks after Veh or TAM treatment, Nestin-Cre^ERT2^::Lin28a^flox/flox^ mice were trained in context A and received a footshock, and freezing was tested 24 hours later in the same context. **(B)** Loss of Lin28a from hippocampal NPCs did not alter freezing of mice during the single-trial contextual fear memory test. (Veh n=13 mice, TAM n=10 mice; two-tailed unpaired t-test, t_21_=0.02071, P=0.8379). **(C)** Experimental paradigm for novel object recognition test. **(D)** Loss of Lin28a from hippocampal NPCs did not impair the novel object recognition of TAM-treated Nestin-Cre^ERT2^::Lin28a^flox/flox^ mice. (Veh n=14 mice, two-tailed paired t-test, t_13_=3.769, **P=0.0023; TAM n=10 mice, two-tailed paired t-test, t_9_=2.908, *P=0.0174). **(E)** Veh-and TAM-treated mice showed similar discrimination index between novel and familiar objects. (Veh n=14 mice, TAM n=10 mice, two-tailed unpaired t-test, t_22_=0.3452, P=0.7332). **(F)** Experimental paradigm for novel location recognition test. **(G)** Loss of Lin28a from hippocampal NPCs impaired the novel location recognition of TAM-treated Nestin-Cre^ERT2^::Lin28a^flox/flox^ mice. (Veh n=14 mice, two-tailed paired t-test, t_13_=5.355, ***P=0.0001; TAM n=12 mice, two-tailed paired t-test, t_11_=1.896, P=0.0845). **(H)** Loss of Lin28a from hippocampal NPCs decreased the discrimination index between novel and familiar locations. (Veh n=14 mice, TAM n=12 mice, two-tailed unpaired t-test, t_24_=2.189, *P=0.0386).

